# Heat-dependent engagement of a buried mitophagy motif in prohibitin 2

**DOI:** 10.64898/2026.10.26.761849

**Authors:** Xinnuo Yang, Yinuo Lin, Zhehua Shao, Xuwen Wang, Binben Wang, Duoduo Zhao, Binchen Liu, Jinliang Nan

**Author notes:** Xinnuo Yang, Yinuo Lin and Zhehua Shao contributed equally as first authors. Corresponding authors: Jinliang Nan and Bingchen Liu.

## Abstract

**Mitophagy receptors are generally switched on by upstream enzymes rather than by the receptor itself. Whether a receptor can instead respond directly to a physical stimulus has not been established. Here we show that a second LC3-interacting region (LIR) in prohibitin 2 (PHB2) is regulated by temperature. It lies in a marginally stable helix that holds its two anchor residues about 5 Å apart, whereas a bound LIR spans about 9 Å. Replica-scaling simulations showed that this helix, but not a matched control from the same protein under identical scaling, samples that geometry; purified PHB2 bound ATG8-family proteins only at 313 K and above. Substitutions designed from this geometry to stabilise (K253A/R255A) or destabilise (I254G) the helix altered binding, mitochondrial recruitment and respiration in PHB2-null cardiomyocytes. Mice carrying the stabilising substitutions in cardiomyocytes sustained greater cardiac injury during heat stroke despite matched thermal load and comparable systemic injury. Motif accessibility may therefore be a regulated variable rather than a fixed property of a fold.**

**Graphical Abstract:** 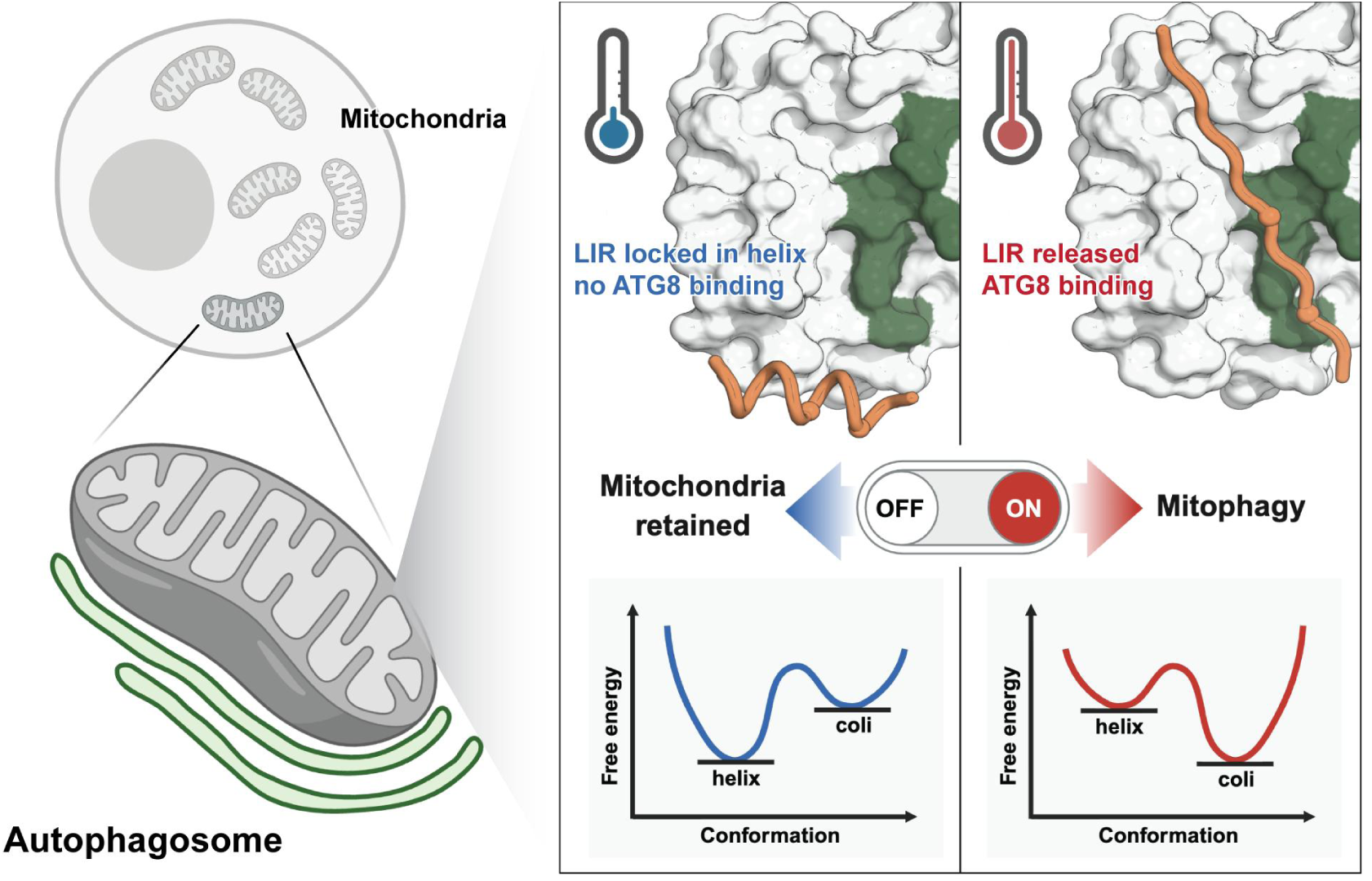

## Introduction

Heat damages mitochondria quickly. Heating inactivates respiratory complexes and depolarises the inner membrane over tens of minutes^1,2^. A cell that must keep producing ATP under these conditions has to clear damaged mitochondria promptly, and the fewer steps that separate the insult from the receptor that acts on it, the sooner clearance can begin. Cardiomyocytes depend on this particularly heavily: they rely almost entirely on oxidative phosphorylation, and the heart is among the first organs to fail in heat stroke^3,4^. Heat-attributable ischaemic heart disease mortality in Asia has risen with the accelerating warming trend, so the cell-level response to heat has a growing clinical stake^5,6^.

Mitochondria can be marked for degradation either by ubiquitination downstream of PINK1 and Parkin, or by membrane-anchored receptors that bind ATG8 proteins directly^7^. In the receptors characterised so far, the receptor is switched on by upstream signalling rather than by the receptor itself. FUNDC1 and BNIP3 are activated by phosphorylation^8,9^, NIX largely by transcriptional upregulation, and others by proteolytic processing^10^. These mechanisms converge on the same endpoint: the receptor’s LC3-interacting region (LIR), an aromatic residue and a downstream hydrophobic residue that insert into two adjacent pockets on an ATG8-family protein, becomes available for binding. But activation of this kind requires an enzymatic cascade to run to completion, so the receptor cannot respond faster than the relay that controls it. Current models therefore place receptor activation downstream of that relay, and do not address whether a receptor can instead respond directly to a physical change in its environment.

The properties of the LIR itself make this gap worth pursuing. The motif is only four residues long and degenerate, so sequence matches run into the thousands across the proteome while only a small fraction are functional^11^. The usual criterion for telling them apart is accessibility: a motif in an intrinsically disordered segment is taken to be usable, one buried in a folded domain to be inert. This treats accessibility as a fixed property of the structure, such that a motif either can or cannot work. The regulatory mechanisms described for LIRs rest on the same premise, acting on top of accessibility rather than on accessibility itself, through phosphorylation of the motif or its flanking residues, or through changes in receptor abundance^12^.

The possibility that a LIR buried within a folded region could be exposed by local unfolding has been raised, but described as largely unexplored^13^. We therefore propose that for some LIRs, accessibility is itself the regulated variable. If the structural element enclosing a motif is only marginally stable, accessibility ceases to be a fixed property and becomes a population that can be shifted. What opens the interaction need not then be an enzyme, but any factor that changes the folding state of that element, of which temperature is the most direct.

Prohibitin 2 (PHB2) offers a system in which to test this. PHB2 is an inner-mitochondrial-membrane mitophagy receptor with a validated LIR at residues 121-124^14^, and it assembles with PHB1 into the ring-shaped complexes that organise the inner membrane^15^. Because it lies in the inner membrane, PHB2 is reached by cytosolic ATG8-family proteins only where the outer membrane has been breached. It also carries a second sequence matching the consensus, YIKL at residues 248-251, which in the solved PHB1-PHB2 structures lies inside an α-helix in both the closed and open states^16,17^, whereas a bound LIR is extended; the two geometries are incompatible. Here we examine the stability of this helix, test whether temperature can regulate PHB2 binding to ATG8-family proteins, and assess the functional consequences in primary cardiomyocytes and in a mouse heat-stroke model.

## Results

### Heat-induced recruitment points to PHB2

Primary neonatal cardiomyocytes held at 315 K accumulated more GABARAPL2 on mitochondria than cells held at 310 K, measured as Mander’s M1 for overlap with TOM20 (difference 0.146, Šídák-corrected 95% CI 0.012 to 0.280; Fig. 1a,b). To distinguish increased delivery from impaired clearance, we repeated the comparison with bafilomycin A1. Bafilomycin raised colocalisation at 315 K as well as at 310 K, so heat had not already saturated the degradative step, and with that step clamped colocalisation remained higher at 315 K (difference 0.230, Šídák-corrected 95% CI 0.096 to 0.364; Fig. 1b). The bafilomycin-sensitive increment was larger at 315 K, but this interaction was not significant (0.084, 95% CI -0.032 to 0.200). Heat therefore increases delivery of GABARAPL2 to mitochondria rather than blocking its removal.

**Figure 1.**
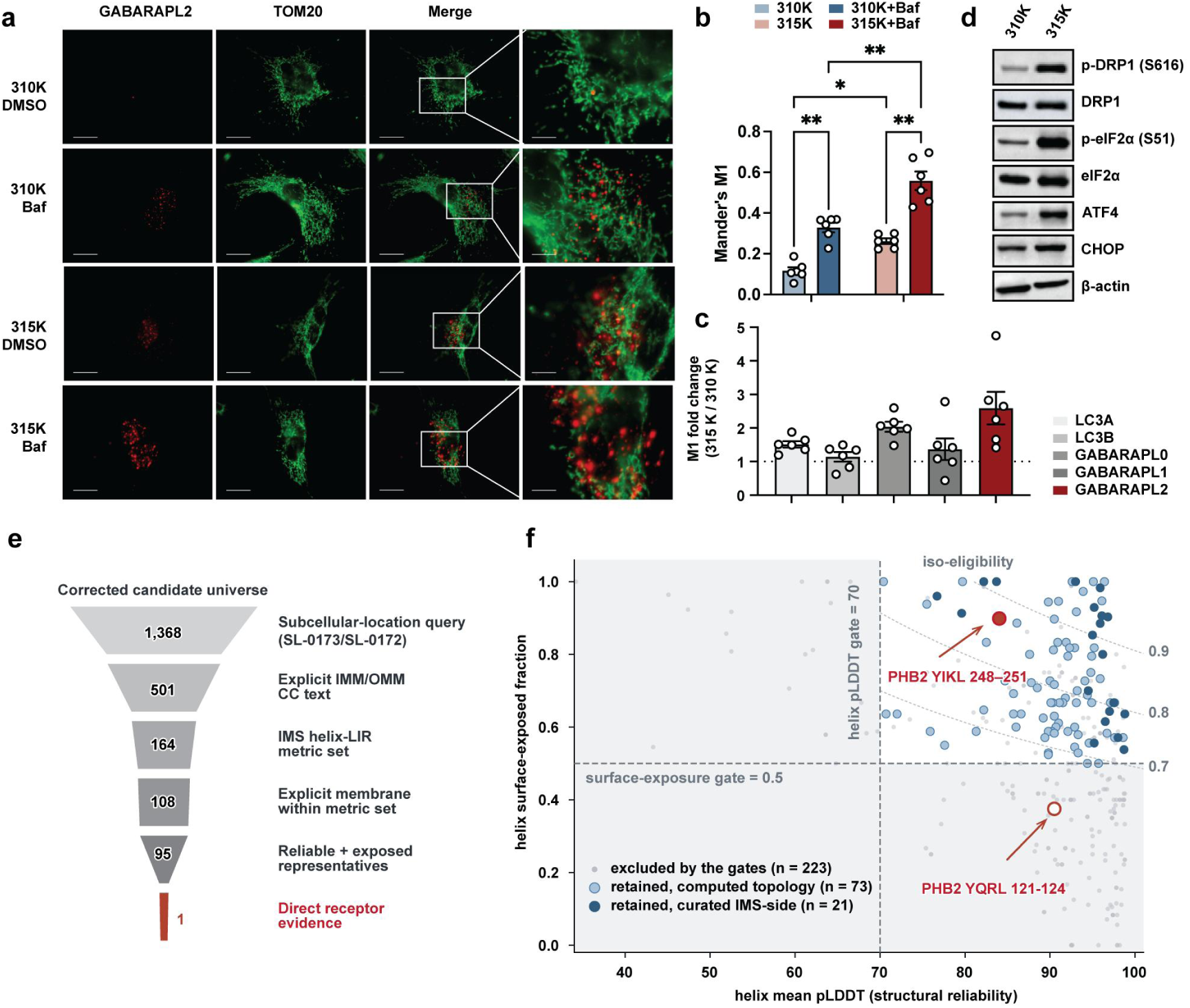
Heat increases GABARAPL2 delivery to mitochondria and a structure-aware screen converges on PHB2. **a,** Immunofluorescence colocalisation of GABARAPL2 (red) with TOM20 (green) in primary neonatal cardiomyocytes held at 310 K or 315 K, each with vehicle (DMSO) or bafilomycin A1. The right-hand column shows the boxed region enlarged. Representative of n = 6 independent isolations. Scale bars, 10 μm (2 μm in enlargements). **b,** Colocalisation in a quantified as Mander’s M1 (Costes automatic thresholds). Bars, mean ± s.e.m.; points, independent cardiomyocyte isolations (n = 6, 20 cells each), with statistics computed on isolation-level means. Repeated-measures two-way ANOVA with both factors matched: temperature F(1,5) = 65.2, P = 0.0005; bafilomycin F(1,5) = 62.3, P = 0.0005; interaction F(1,5) = 3.44, P = 0.12. All six pairwise comparisons of cell means were made with a common error term and Šídák correction; the four marked are bafilomycin at 310 K, P = 0.0070, and at 315 K, P = 0.0015, and 315 K versus 310 K in vehicle, P = 0.036, and with bafilomycin, P = 0.0049. *P < 0.05, **P < 0.01. **c,** Colocalisation of five ATG8-family proteins with TOM20 at 315 K, expressed as the fold change over the matched 310 K control for each paralogue. The dashed line marks no change. Bars, geometric mean with 95% confidence interval; points, independent isolations (n = 6). Ratios were tested on a log scale against unity and corrected across the five paralogues by the Holm method: GABARAPL0 2.01-fold, *P* = 0.0012; LC3A 1.50-fold, *P* = 0.0055; GABARAPL2 2.39-fold, *P* = 0.013; GABARAPL1 1.18-fold, *P* = 1.0; LC3B 1.09-fold, *P* = 1.0. GABARAPL0 and GABARAPL2 were not distinguishable from each other (1.19-fold, 95% CI 0.65 to 2.18, P = 0.49). **d,** Immunoblots of cardiomyocytes held at 310 K or 315 K, probed for p-DRP1 (S616), DRP1, p-eIF2α (S51), eIF2α, ATF4 and CHOP, with β-actin as loading control. Representative of n = 3 independent isolations. **e,** Candidate funnel. A subcellular-location query returned 1,368 mitochondrial proteins; requiring an explicit inner- or outer-membrane annotation reduced this to 501, of which 164 carried a candidate LIR falling within a helix and 108 retained an explicit membrane annotation on review. Applying the two prespecified structural gates and keeping one representative site per protein left 95 proteins, of which one (PHB2) also had established mitophagy-receptor evidence. **f,** The eligibility plane underlying the final step of e: mean pLDDT of the helix carrying the candidate LIR against the solvent-exposed fraction of that helix. Plotted are the 223 excluded motifs and the 95 retained proteins, each represented by one site. Eligibility is √(exposure × pLDDT/100); dashed lines are the prespecified gates (pLDDT ≥ 70; exposure ≥ 0.5) and dotted lines are iso-eligibility contours. Grey points were excluded by the gates (n = 223); light and dark blue are retained candidates assigned by computed topology (n = 73) and by curated IMS-side assignment (n = 21), with PHB2 YIKL shown separately in red. Both PHB2 LIRs are marked: YIKL 248-251 (filled) passes both gates, whereas YQRL 121-124 (open) is excluded by insufficient exposure.

Comparing all five human ATG8-family proteins, each normalised to its own 310 K control so that differences in antibody affinity cancel, GABARAPL0 (also known as GABARAP) and GABARAPL2 showed the largest increases (2.0-fold, 95% CI 1.7 to 2.4, and 2.4-fold, 95% CI 1.5 to 3.8) and were not distinguishable from each other (*P* = 0.49). LC3A increased less (1.5-fold, 95% CI 1.3 to 1.8; Fig. 1c and Extended Data Fig. 1a–d). LC3B and GABARAPL1 did not increase detectably (1.1-fold, 95% CI 0.8 to 1.6, and 1.2-fold, 95% CI 0.6 to 2.2); these intervals include unity, and we do not read them as evidence of no change.

Heat also engaged the integrated stress response and altered mitochondrial dynamics: immunoblots showed higher phosphorylation of eIF2α at Ser51 and of DRP1 at Ser616 and higher ATF4 and CHOP (Fig. 1d), and among mitochondrial markers the clearest change was a reduction in SOD2 (Extended Data Fig. 1e). These responses indicate that the exposure produced a bona fide stress state.

We then asked which mitochondrial protein could present a LIR under these conditions. A structure-aware screen began with 1,368 proteins from a mitochondrial location query, retained 501 with an explicit inner- or outer-membrane annotation, 164 carrying a candidate LIR falling within a helix, and 108 retaining an explicit membrane annotation on review. Two structural gates, mean helix pLDDT of at least 70 and surface-exposed fraction of at least 0.50, left 95 representative proteins, of which PHB2 was the only one with established mitophagy-receptor evidence (Fig. 1e).

PHB2 carries two LIR candidates, and the gates separated them (Fig. 1f). The validated YQRL motif at residues 121–124 was predicted to be buried (exposed fraction 0.375) and was excluded, whereas YIKL at residues 248–251 was exposed (0.900) and retained. The helix carrying YIKL was among the least confident retained by the pLDDT gate (84.2, the 15th percentile of the 95 retained) while being more exposed than most (0.900, the 78th percentile), an unusual combination. The motif itself is unremarkable by sequence: its iLIR xLIR score of 11 lies one point below the median of the human distribution (median 12, range 4–26) (Extended Data Fig. 2a). What distinguishes it is structural context. PHB2 ranked 25th of 95 and held that rank at every one of nine gate settings tested, so its retention did not depend on a particular threshold (Extended Data Fig. 2b). An evidence matrix makes explicit that receptor status is established at the protein level while YIKL remains a predicted site (Extended Data Fig. 2c).

### A marginally stable helix buries the LIR

Residues 246-257 (PGYIKLRKIRAA) are helical in the available PHB1-PHB2 cryo-electron microscopy structures. Across 33 PHB2 chains from PDB entries 9O9Z, 9OA0 and 9UNL, covering both the closed and open states of the complex, the Cα atoms of the two LIR anchors, Tyr248 and Leu251, lie between 4.94 and 5.27 Å (Fig. 2a and Supplementary Table 1). A LIR bound in an ATG8 groove spans considerably more: 8.99 Å in the GABARAPL2-IRGQ complex^18^, and 8.63-9.63 Å across seven human ATG8-LIR complexes (Supplementary Table 2). No rigid α-helix can present both anchors at that separation, so for PHB2 to be read as a LIR the helix must first come apart.

**Figure 2.**
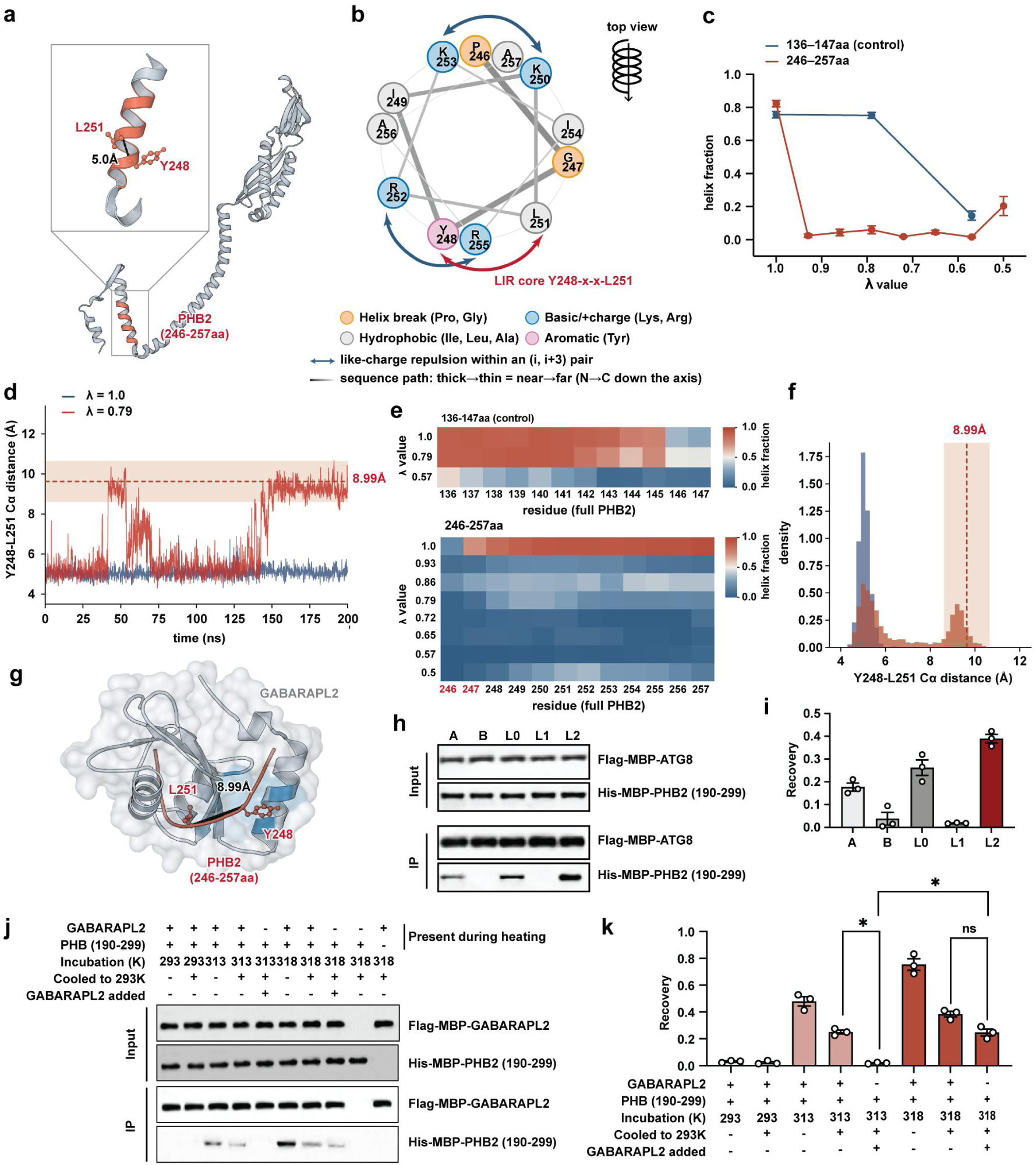
A marginally stable helix holds the PHB2 LIR anchors too close for the ATG8 groove, and binding requires elevated temperature. **a,** AlphaFold model of full-length human PHB2 (UniProt Q99623) with residues 246-257 in red. The inset enlarges this segment; the Cα atoms of Tyr248 and Leu251 are 5.0 Å apart in this model. **b,** Helical wheel of PHB2 246-257 viewed down the helix axis. Colours distinguish helix-breaking residues (Pro, Gly), basic residues, hydrophobic residues and the aromatic residue; double-headed arrows mark like-charge repulsion within (i, i+3) pairs, and the grey path runs from thick to thin along the sequence from N to C terminus. The LIR core Y248-x-x-L251 is marked in red. **c,** Helical fraction of the target segment 246-257 (red) and the control segment 136-147 from the same protein (blue) across Hamiltonian scaling factors λ. Each λ was run as a single trajectory; points are the mean of ten equal time blocks spanning the final 100 ns, with s.e.m. over those blocks. **d,** Cα-Cα separation of Tyr248 and Leu251 over 200 ns at λ = 1.0 (blue) and λ = 0.79 (red). The shaded band is 8.99 ± 1 Å, the separation in the GABARAPL2-IRGQ complex (PDB 8Q6Q) with a 1 Å tolerance. **e,** Per-residue helical fraction at each λ for the control segment 136-147 (top) and the target segment 246-257 (bottom). Pro246-Gly247 is marked in red. **f,** Full-trajectory distributions of the two traces in **d. g,** A frame from the λ = 0.79 trajectory (red), in which the Tyr248-Leu251 separation is 9.63 Å, superposed on the bound-LIR position in the GABARAPL2 groove (grey surface), placing Tyr248 and Leu251 in the two hydrophobic pockets. The labelled 8.99 Å is the anchor separation required by the groove, measured in the GABARAPL2-IRGQ complex, not the separation in the displayed frame. h, *In vitro* co-immunoprecipitation with purified proteins incubated at 318 K. Flag-MBP-tagged ATG8-family proteins (A, LC3A; B, LC3B; L0, GABARAPL0; L1, GABARAPL1; L2, GABARAPL2) were used as bait and incubated with His-MBP-PHB2(190-299). Representative of n = 3 independent experiments. **i,** Densitometric quantification of h. Recovery is (IP PHB2 / input PHB2) ÷ (IP bait / input bait). Bars, mean ± s.e.m.; points, independent experiments (n = 3). j, *In vitro* co-immunoprecipitation testing whether the temperature-dependent change is reversible. PHB2(190-299) was incubated with or without Flag-MBP-GABARAPL2 at 293, 313 or 318 K, returned to 293 K where indicated, and in two conditions the receptor was added only after cooling. The condition matrix below the panel gives the composition of each lane. The last two lanes are specificity controls at 318 K, omitting the receptor and the PHB2 fragment respectively. Representative of n = 3 independent experiments. **k,** Densitometric quantification of the first eight lanes of **j**, calculated as in **i.** Bars, mean ± s.e.m.; points, independent experiments (n = 3). The dashed line marks the background level, taken as the mean of the two 293 K conditions. Repeated-measures one-way ANOVA with Geisser-Greenhouse correction; five preselected comparisons were made with the Holm-Šídák method. The three marked are lane 5 versus lane 4, *P* = 0.016; lane 8 versus lane 5, *P* = 0.045; and lane 8 versus lane 7, *P* = 0.18. \**P* < 0.05; ns, not significant.

Several features of the segment predict low helical stability. Viewed down its axis it carries no acidic residue and four basic ones, which cluster on a single face and form two like-charge (i, i+3) pairs (Fig. 2b); its hydrophobic moment is low (μH = 0.263, 68% that of the control helix at residues 136-147)^19^; and it begins with the helix-breaking pair Pro246-Gly247. The segment is nonetheless identical in human, mouse and rat, and in 35 of the 36 vertebrate orthologues examined; the exception, Xenopus tropicalis, substitutes Ile249Leu and Lys253Arg, so that Tyr248, Leu251 and the basic face are conserved throughout (Supplementary Table 3).

We tested this in simulation by replica-scaling molecular dynamics^20^, in which λ sets an effective temperature for the scaled segment far above the physiological range, to accelerate transitions that are otherwise too slow to sample. A control helix from the same protein, residues 136-147, stayed folded for the full 200 ns at λ = 1.0 and λ = 0.79, whereas the 246-257 segment stayed folded only at λ = 1.0 and lost its helix at every reduced λ tested (Fig. 2c and Extended Data Fig. 3a,b). The comparison this supports is the matched one: at λ = 0.79 the target segment unfolds while the control, in the same protein and under identical scaling, does not. Unfolding began at Pro246-Gly247, where helix occupancy was 0.25 and 0.32 over the first 30 ns against 0.89-0.95 for Arg252-Ala257 (Fig. 2e). Solvent exposure rose over most of the target segment but changed little across the control (Extended Data Fig. 3c,d). Once the helix opened, the anchors reached the required spacing: 54.0% of frames in the final 100 ns at λ = 0.79 fell within 8.99 ± 1 Å, against 0% at λ = 1.0 (Fig. 2d,f). Superposing one such frame on the bound-LIR position in the GABARAPL2 groove places both anchors in the two hydrophobic pockets without steric conflict (Fig. 2g). At λ = 0.79 the required geometry is therefore reachable by this segment and not by the matched control.

We then simulated the extended PHB2 LIR bound to each of the five ATG8-family proteins, to ask whether that conformation is stable in a groove rather than merely able to reach one. The peptide was retained in four of five systems, with the interface carried by a small number of discrete contacts in every system; GABARAPL2 showed the closest agreement between replicates, and the one trajectory that dissociated was GABARAPL1 (Extended Data Fig. 4). These simulations test whether a modelled complex holds together rather than how the receptors rank, so we measured binding directly. Purified Flag-MBP-tagged receptors incubated with His-MBP-PHB2(190-299) at 318 K recovered PHB2, most in the GABARAPL2 and GABARAPL0 lanes (recovery 0.390 and 0.262), although these two were not distinguishable (P = 0.114; n = 3 independent experiments; Fig. 2h,i). Across a temperature series, no receptor recovered PHB2 at 293-308 K; recovery appeared at 313 K for GABARAPL2, GABARAPL0 and LC3A, and only at 318 K for LC3B and GABARAPL1, increasing at 318 K in every case (Extended Data Fig. 5). These reactions contain only the two purified proteins, so the temperature dependence is a property of the pair rather than of the cellular environment, and it places the threshold for detectable binding between 308 and 313 K.

To ask whether the change persists after cooling, we heated PHB2(190-299) on its own, returned it to 293 K, and only then added GABARAPL2. If heating leaves a lasting change in PHB2, binding should still be recovered under these conditions; if it does not, the receptor has nothing to engage. After 318 K, PHB2 was still recovered: more than in the corresponding 313 K condition (0.228, *P* = 0.045), and not detectably less than when the receptor had been present throughout the incubation (-0.137, *P* = 0.18; Fig. 2j,k). After 313 K, no recovery above background was detected, and recovery was lower than when the receptor had been present during the same heating step (-0.231, *P* = 0.016). Heating to 318 K therefore leaves PHB2 in a state that still engages GABARAPL2 after the sample has returned to 293 K.

### Substitutions that alter helix stability

The wheel analysis specified where to mutate. Replacing Lys253 and Arg255 with alanine removes one partner from each (i, i+3) pair and abolishes both same-face repulsions, while alanine is the strongest helix former (ΔΔG +0.47 kcal mol⁻¹ from propensity alone)^21^; Tyr248 and Leu251 are untouched, so these substitutions are designed to stabilise the helix rather than to delete the motif. I254G was designed to act in the opposite direction. An AlphaFold3 comparison^22^ across seven candidate substitutions showed that neither K253A/R255A nor I254G produced a resolved change in the folded state, whereas A256P and R252P distorted the flanking geometry and were discarded (Extended Data Fig. 6). We therefore took forward only substitutions predicted to leave the folded structure unchanged, so that a functional difference is less readily explained by a damaged fold.

We expressed wild-type PHB2, K253A/R255A or I254G in PHB2-knockout cardiomyocytes, so that each variant is the cell’s only PHB2. Immunoblots of GABARAPL2 pull-downs recovered PHB2 after heat in wild-type cells, with a fainter band for K253A/R255A and a stronger band for I254G (Fig. 3a). The three variants were expressed at comparable levels, PHB1 was unchanged, and both variants retained PHB1-PHB2 complex assembly, which was absent in knockout cells (Fig. 3b), excluding gross disruption of the prohibitin complex. Mitochondria-associated GABARAPL2 rose with heat in wild-type and I254G cells (+0.202, Šídák-corrected 95% CI +0.104 to +0.300, and +0.134, +0.036 to +0.231) but not detectably in K253A/R255A cells (+0.072, -0.026 to +0.170), and the effect of genotype depended on temperature (interaction F(2,10) = 6.44, *P* = 0.016; Fig. 3c,d). After heat, K253A/R255A retained less GABARAPL2 than either wild type (-0.150, -0.248 to -0.052) or I254G (-0.235, -0.332 to -0.137), while I254G did not differ detectably from wild type (+0.085, -0.013 to +0.182). After heat, the strongest bands for LONP1, CHOP and phosphorylated AMPK, and the weakest for SOD2 and MFN2, were in K253A/R255A cells, with I254G cells showing the converse (Fig. 3e,f).

**Figure 3.**
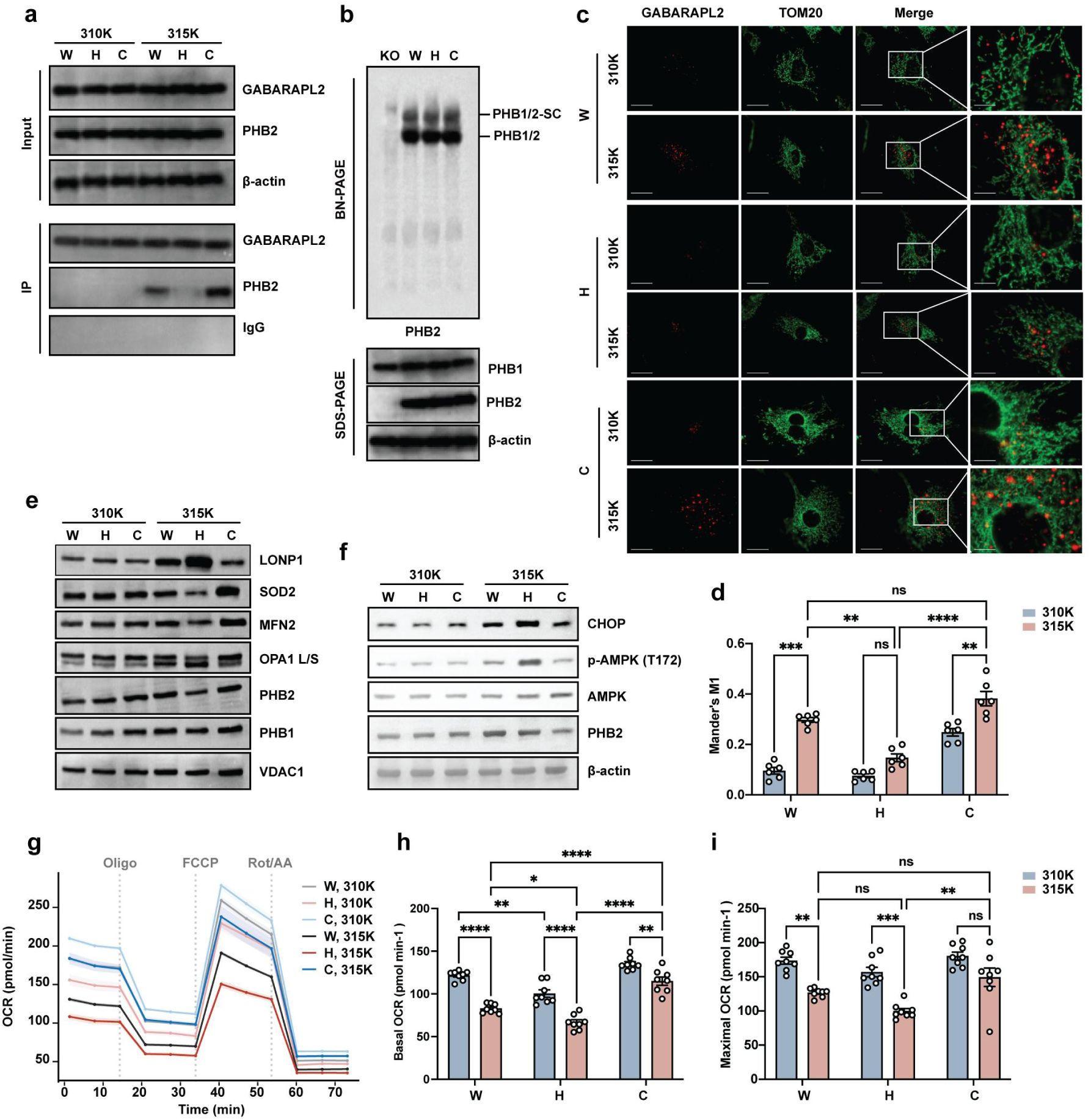
Helix-stabilising and helix-destabilising substitutions in PHB2-knockout cardiomyocytes. Throughout, PHB2-knockout cardiomyocytes were rescued with wild-type PHB2 (W), K253A/R255A (H) or I254G (C) and held at 310 K or 315 K. **a,** Co-immunoprecipitation of PHB2 with GABARAPL2. Input and immunoprecipitate were probed for GABARAPL2 and PHB2, with β-actin as a loading control for the input and an IgG immunoprecipitation as a specificity control. **b,** Blue-native PAGE of the same lines and of PHB2-knockout cells (KO), probed for PHB2, resolving the prohibitin complex (PHB1/2) and its supercomplex (PHB1/2-SC). Lower panels, SDS-PAGE of the same lysates probed for PHB1, PHB2 and β-actin. **c, I**mmunofluorescence colocalisation of GABARAPL2 (red) with TOM20 (green). The fourth column shows the boxed region enlarged. Representative of n = 6 independent isolations. Scale bars, 10 μm (2 μm in enlargements). **d,** Quantification of **c** as Mander’s M1 with Costes automatic thresholds. Bars are mean ± s.e.m.; points are independent isolations (n = 6, 20 cells each), with statistics computed on isolation-level means. Repeated-measures two-way ANOVA with matching on both factors: genotype F(2,10) = 103.4, *P* < 0.0001; temperature F(1,5) = 82.1, *P* = 0.0003; interaction F(2,10) = 6.44, *P* = 0.016. All fifteen pairwise comparisons of cell means were made with Šídák’s test; six are marked in the panel. \*\**P* < 0.01, \*\*\**P* < 0.001, \*\*\*\**P* < 0.0001; ns, not significant. **e,** Immunoblots probed for LONP1, SOD2, MFN2, the long and short forms of OPA1, PHB2 and PHB1, with VDAC1 as a mitochondrial loading control. **f,** Immunoblots probed for CHOP, phosphorylated AMPK (Thr172), total AMPK and PHB2, with β-actin as a loading control. **g,** Oxygen consumption rate through a mitochondrial stress test; dotted lines mark the injection of oligomycin, FCCP, and rotenone with antimycin A. Lines are mean, shading s.e.m. (n = 8 independent isolations). **h,i,** Basal (**h**) and maximal (**i**) oxygen consumption rate derived from **g.** Bars are mean ± s.e.m.; points are independent isolations (n = 8). Repeated-measures two-way ANOVA with matching on both factors: basal, genotype F(2,14) = 103.1, *P* < 0.0001, temperature F(1,7) = 369.3, *P* < 0.0001, interaction F(2,14) = 4.91, *P* = 0.024; maximal, genotype F(2,14) = 13.2, *P* = 0.0006, temperature F(1,7) = 112.4, *P* < 0.0001, interaction F(2,14) = 1.62, *P* = 0.23. All fifteen pairwise comparisons of cell means were made with Tukey’s test; those marked are shown in the panels. In **a**, **b**, **e** and **f**, blots are shown as representative of n = 3 independent isolations.

Respiration tracked the direction of the designed change. Basal oxygen consumption was 100.4 ± 4.4 pmol min⁻¹ for K253A/R255A, 121.7 ± 2.0 for wild type and 135.3 ± 2.6 for I254G at 310 K, and 66.6 ± 3.1, 82.9 ± 1.9 and 114.9 ± 4.8 after heat (mean ± s.e.m., n = 8; Fig. 3g,h). Heat lowered basal respiration in all three lines and the effect of genotype depended on temperature (interaction F(2,14) = 4.91, *P* = 0.024); after heat, K253A/R255A remained below wild type (-16.3 pmol min⁻¹, Tukey-corrected 95% CI -30.4 to -2.1) and I254G above it (+32.0, +17.9 to +46.2). Maximal respiration fell with heat in wild-type and K253A/R255A cells but not detectably in I254G cells (-31.1, -65.0 to +2.9), and after heat K253A/R255A remained below I254G (-49.6, -83.5 to -15.6); genotype and temperature did not interact for this measure (F(2,14) = 1.62, *P* = 0.23; Fig. 3i). K253A/R255A lowered basal respiration already at 310 K, before any heat (-21.3, -35.5 to -7.2). If the helix were fully folded at rest, a substitution that stabilises it further would have no effect; the observed difference therefore indicates that part of the wild-type population is open even at 310 K.

### A knock-in allele worsens cardiac injury

We introduced K253A/R255A into cardiomyocytes in vivo, crossing a conditional knock-in allele to constitutive Myh6-Cre and comparing with Cre-negative floxed littermates. PHB2 was recovered with GABARAPL2 from control hearts after heat stroke, but not detectably from knock-in hearts (Fig. 4a).

**Figure 4.**
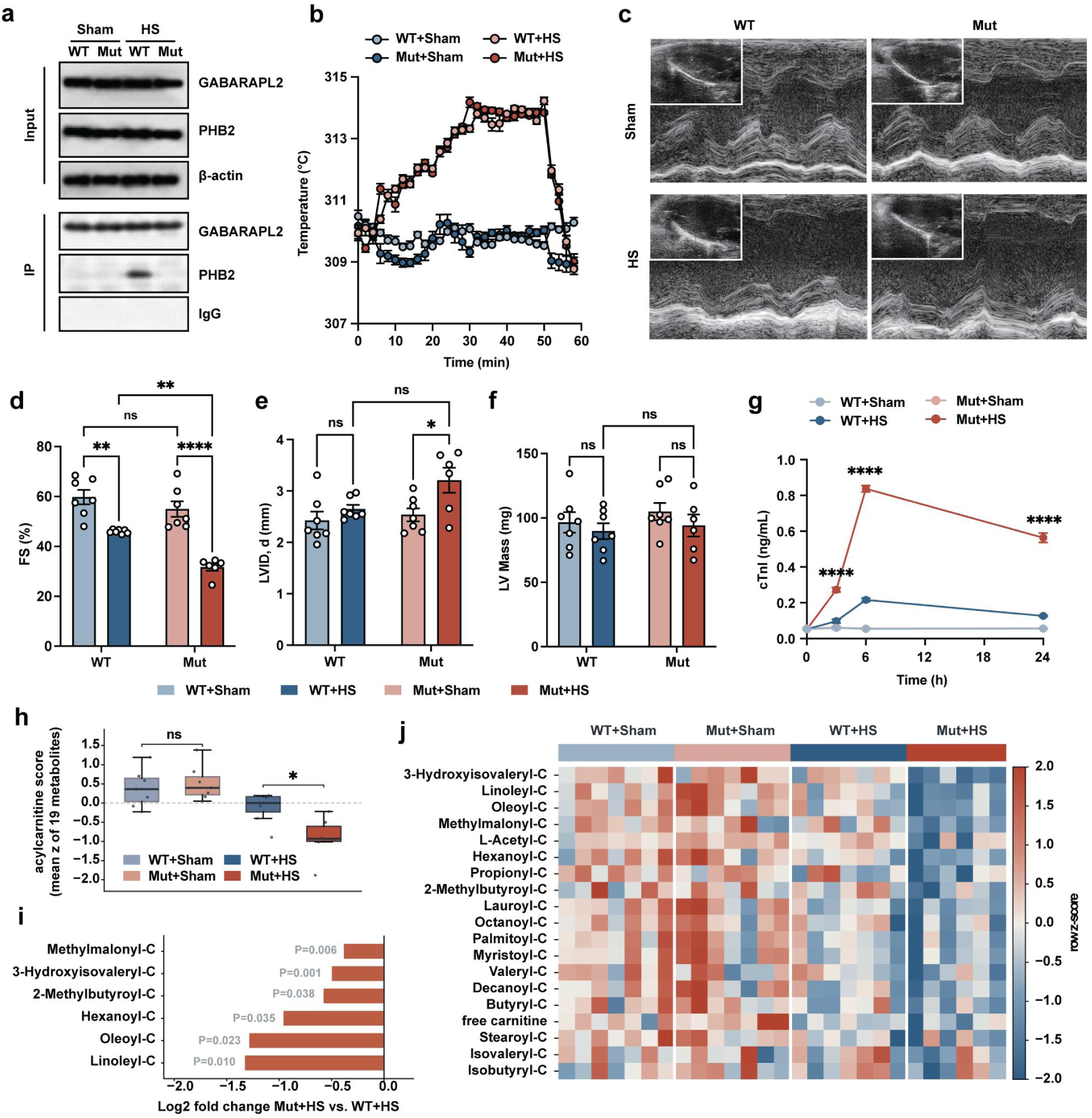
Cardiomyocyte-specific K253A/R255A knock-in mice in a heat-stroke model. Throughout, mice carrying a conditional K253A/R255A knock-in allele crossed to Myh6-Cre (Mut) are compared with Cre-negative floxed littermates (WT), each subjected to heat stroke (HS) or a sham procedure. n = 7 per group except Mut+HS, in which one animal died after the temperature recording and before endpoint sampling (n = 6 for **c-j**). **a,** Co-immunoprecipitation of PHB2 with GABARAPL2 from heart lysates. Input and immunoprecipitate were probed for GABARAPL2 and PHB2, with β-actin as a loading control for the input and an IgG immunoprecipitation as a specificity control. Representative of n = 3 independent experiments. **b,** Core temperature recorded every 2 min for 58 min (n = 7 per group). Points are mean, error bars s.e.m. The two heat-stroke groups did not differ in peak core temperature (difference -0.08 K, 95% CI -0.34 to 0.18, P = 0.50), in mean core temperature across the recording (-0.02 K, 95% CI -0.16 to 0.13, P = 0.81) or in time above 313 K (0.57 min, 95% CI -2.62 to 3.77, P = 0.69; two-tailed Welch’s t-test). **c,** Representative M-mode echocardiograms; insets show the B-mode image from which the M-mode line was taken. **d-f,** M-mode fractional shortening (**d**), left-ventricular internal diameter at end-diastole (**e**) and left-ventricular mass (**f**). Bars are mean ± s.e.m.; points are individual mice. Two-way ANOVA: fractional shortening, genotype F(1,23) = 2.28, *P* = 0.14, surgery F(1,23) = 18.8, *P* = 0.0002, interaction F(1,23) = 4.27, *P* = 0.050; internal diameter, interaction F(1,23) = 1.94, *P* = 0.18; left-ventricular mass, interaction F(1,23) = 0.08, *P* = 0.79. All six pairwise comparisons of the four cell means were made with Tukey’s test; those marked are shown in the panels. \**P* < 0.05, \*\**P* < 0.01, \*\*\*\**P* < 0.0001; ns, not significant. **g,** Serum cardiac troponin I before and 3, 6 and 24 h after the procedure. Points are mean, error bars s.e.m. The lower limit of quantification of the assay was 0.05 ng mL⁻¹ and values below it were assigned that limit; because half of the sham values were at this limit, statistical comparison was restricted to the two heat-stroke groups. Mixed-effects model with matching by time: genotype F(1,11) = 1112, *P* < 0.0001; time F(3,33) = 488, *P* < 0.0001; genotype × time F(3,33) = 242, *P* < 0.0001. Asterisks compare Mut+HS with WT+HS at each time point (Šídák-corrected across the four time points); the two groups did not differ before the procedure (*P* = 1.00). The sham groups were not tested. \*\*\*\**P* < 0.0001. **h,** Composite acylcarnitine score, the mean row z-score of the 19 cardiac carnitine species measured (free carnitine and 18 acylcarnitines). Boxes show the median and interquartile range, whiskers the full range, and points individual mice. Two-way ANOVA genotype × surgery interaction, *P* = 0.019. \**P* < 0.05; ns, not significant. **i,** Log2 fold change in knock-in relative to control hearts after heat stroke, for the six of the nineteen species with uncorrected *P* < 0.05 (Welch’s t-test). *P* values are not corrected for multiple comparisons. **j,** Row z-scores for the 19 carnitine species used in **h**, in every animal. All species of this chemical class that were measured are shown; none were selected on the data. Metabolites are ordered by their mean z-score in the Mut+HS group, most depleted at the top.

Both genotypes experienced the same thermal load. Peak core temperature did not differ between genotypes (314.55 ± 0.05 K and 314.47 ± 0.10 K; difference -0.08 K, 95% CI -0.33 to +0.17), nor did mean core temperature across the recording (-0.02 K, -0.16 to +0.13) or time above 313 K (26.3 ± 1.3 min and 26.9 ± 0.6 min; difference +0.6 min, -2.5 to +3.6), while both sham groups remained normothermic (n = 7 per group; Fig. 4b and Extended Data Fig. 7a). Serum IL-6, TNF-α and IL-1β and the liver and kidney injury markers ALT, AST, creatinine and urea all rose after heat stroke, and none differed between genotypes when the seven comparisons were corrected together (Extended Data Fig. 7b).

Against that matched background the cardiac phenotype was clear. One knock-in heat-stroke animal died after the temperature recording and before endpoint sampling, leaving n = 6 in that group; its exclusion biases the comparison towards the null. In sham animals the two genotypes were indistinguishable in fractional shortening (difference -4.8, Tukey-corrected 95% CI -13.6 to +4.0), left-ventricular internal diameter (+0.11 mm, -0.51 to +0.73) and left-ventricular mass (+8.3 mg, -19.9 to +36.6), so the allele carries no detectable baseline cardiac cost despite being expressed from development. Heat stroke lowered fractional shortening in both genotypes, by 13.8 (-22.6 to -5.0) in controls and by 23.3 (-32.5 to -14.1) in knock-in mice, and after heat stroke it was substantially lower in knock-in hearts (-14.3, -23.5 to -5.1, *P* = 0.0014), although the genotype × surgery interaction was borderline (*P* = 0.050; Fig. 4c,d). Left-ventricular internal diameter increased in knock-in mice after heat stroke (+0.67 mm, +0.03 to +1.31, *P* = 0.039; Fig. 4e), while left-ventricular mass did not differ detectably between any groups (Fig. 4f). Serum cardiac troponin I was at the assay floor in both sham groups throughout, and rose after heat stroke to a peak at 6 h in both genotypes. The peak was 3.9-fold higher in knock-in mice (0.837 ± 0.017 versus 0.217 ± 0.010 ng mL⁻¹; difference +0.62, 95% CI +0.57 to +0.67), with the two genotypes indistinguishable before heat stroke and still separated at 24 h (Fig. 4g).

Targeted cardiac metabolomics showed the same conditional pattern. Because knock-in hearts also sustained more myocyte damage, some loss of small metabolites from injured cells cannot be excluded; the sixfold difference in effect size between acylcarnitines and amino acids indicates that the change is at least partly selective. A composite score over the 19 cardiac carnitine species did not differ between the two sham groups (*P* = 0.61) and fell after heat stroke, further in knock-in mice (*P* = 0.018), giving a genotype × surgery interaction of *P* = 0.019 (Fig. 4h). The difference was carried by long- and branched-chain acylcarnitines, including linoleyl- and oleoylcarnitine (Fig. 4i), and is shown for all 19 species and every animal, with no selection of metabolites on their statistics (Fig. 4j).

## Discussion

This work identifies a mode of receptor activation that had not previously been tested. The helix enclosing YIKL lost structure under reduced Hamiltonian scaling while a matched control helix from the same protein did not at the same λ; once it unfolded, the anchor residues reached the separation the receptor groove requires; and a substitution designed to stabilise the helix weakened binding while one designed to destabilise it strengthened binding. A contrary result in any of these three tests would have defeated the conformational-gating explanation, and all three point the same way. Binding remained temperature-dependent in a reaction containing only the two purified proteins, placing the transition in the pair itself rather than requiring chaperones or post-translational modification. The possibility that a buried motif could be exposed by local unfolding had been raised but not explored; here we test it experimentally and identify temperature as the variable that the interaction depends on.

The gate is graded rather than switch-like. At 310 K, before any heat, the three variants already differed in basal respiration, ordered by the direction of the designed change; a switch-like gate predicts no difference under this condition. The population model also accounts for an in vivo result that appears contradictory at first: mice carrying the stabilising allele had normal baseline cardiac function and normal cardiac morphology, yet sustained markedly greater injury during heat stroke. The fraction open at rest is small, so removing it costs little; heat should raise that fraction, and it is then that the inability to raise it becomes costly. The substitution was designed backwards from the helical-wheel geometry, removing both same-face like-charge pairs without touching Tyr248 or Leu251, and it preserves PHB1-PHB2 complex assembly, so its phenotype cannot be attributed to loss of the motif or to disassembly of the complex.

This result changes how buried motifs should be judged. Sequence matches to the LIR consensus run into the thousands across the proteome, and they are conventionally sorted into usable and inert on the basis of accessibility, which is treated as a fixed property of the structure. If accessibility can move with the stability of the element enclosing it, that dichotomy no longer holds: some of the matches judged inert because they are buried may simply not yet have been opened, and identifying them calls not for better sequence scoring but for an assessment of local structural stability. PHB2 residues 246-257 are identical in 35 of the 36 vertebrate orthologues examined, which suggests that this marginal stability is itself a selected property. At the level of the receptor, GABARAPL1 was the weakest in all three measurements, cell imaging, purified binding and simulation stability, while GABARAPL2 was among the strongest in each, although it was not separable from GABARAPL0 either by imaging (P = 0.49) or by purified binding (P = 0.114). The preference is not absolute, since binding of all five ATG8 proteins increased with temperature, consistent with the established view that LIR subfamily selectivity depends on context beyond the four-residue core^23^.

The conclusions have a defined scope. That temperature changes the conformational state of this segment is supported jointly by three independent lines of evidence: geometry, simulation and purified binding. In replica scaling, λ sets an effective temperature for the scaled segment well above the physiological range, so these simulations report the lability of this segment relative to other helices in the same protein rather than a melting temperature. The change produced at 318 K did not fully reverse when the sample was returned to 293 K, which a simple shift in folding equilibrium would not predict; the peak core temperature reached in vivo, 314.5 K, lies below that point, but whether an additional irreversible process contributes at the highest temperatures has not been resolved. In cells and in vivo, helix stability tracked respiratory capacity and the severity of heat-stroke cardiac injury; the portion of this that proceeds through mitophagy has not been separated from the portion that may proceed through other functions of PHB2. The fall in cardiac acylcarnitines in mutant mice accompanied myocyte injury, and the causal order between them is not established, although the roughly sixfold difference in effect size between acylcarnitines and amino acids indicates that the change is at least partly selective.

Temperature is only one way to move a folding equilibrium. Membrane lipid composition, ionic strength, mechanical tension and ligand binding also alter local stability, and none of these is constant at the inner mitochondrial membrane. A helix that can be opened by temperature can in principle be opened by any of them. Conformational gating may therefore not be specific to heat stress, but a more general mode of receptor regulation in which the local stability of the receptor itself is the receiving end of the signal, with no upstream enzyme required to translate an external state into a chemical modification.

## Acknowledgements

We thank Ms. Yingying Huang and Ms. Yanwei Li from the Core Facilities, Zhejiang University School of Medicine for their technical support.

## Author Contributions Statement

J.N., X.Z. and Z.S. conceived, designed and supervised the overall project. J.N. and Z.S. performed most experiments and analysed data. Y.L, X.Y. and D.Z. contributed to in vitro experiments, mouse model generation, and biochemical analyses. J.N., X.W. and B.W. performed molecular dynamics simulations and structural analyses. D.Z. assisted with mitochondrial isolation, respirometry, and imaging studies. J.N. and Z.S. wrote the manuscript with assistance of B.L.

## Methods

Temperatures that define an experimental condition are given in kelvin: 310 K and 315 K denote incubator set points of 37 °C and 42 °C, the in vitro binding series spans 293-318 K, and animal housing, chamber and core temperatures are likewise given in kelvin. Routine procedural temperatures, such as centrifugation and incubation steps, are given in degrees Celsius.

### Primary neonatal cardiomyocyte isolation and heat treatment

Primary neonatal mouse cardiomyocytes were isolated from postnatal day 1-3 Phb2^flox/flox^ mice of both sexes. Ventricles were excised, minced into approximately 1-mm³ fragments and subjected to repeated enzymatic digestion with 0.125% trypsin and 0.1% collagenase II at 37 °C. The dissociated cells were collected, filtered through a 70-µm cell strainer and pre-plated for 90 min to reduce fibroblast contamination. Non-adherent cardiomyocytes were seeded at approximately 1 × 10^5^ cells cm^-2^ on gelatin-coated culture plates in DMEM/F12 supplemented with 10% fetal bovine serum, 1% penicillin-streptomycin and 0.1 mM 5-bromo-2’-deoxyuridine. After 24 h, the medium was replaced with DMEM/F12 containing 2% fetal bovine serum. Cardiomyocyte purity was confirmed by cardiac troponin T immunostaining and routinely exceeded 90%.

For heat-stress experiments, cardiomyocytes were maintained at 310 K or transferred to a pre-equilibrated humidified incubator at 315 K with 5% CO₂ for 2 h. Incubator temperature was verified with a calibrated thermometer at the culture-vessel level before each experiment. Cells were collected immediately after treatment for imaging, immunoblotting, co-immunoprecipitation and mitochondrial-function assays. Where indicated, cells were pretreated with 100 nM bafilomycin A1 for 2 h, and the inhibitor was maintained throughout the temperature exposure. An equivalent volume of dimethyl sulfoxide was added to vehicle-control cultures. Bafilomycin A1 was used to test whether mitochondria-associated ATG8 signal continues to accumulate when lysosomal degradation is blocked. The temperature × bafilomycin interaction, rather than either treatment alone, was used as the basis for inferring a change in flux through the pathway.

### PHB2 depletion and variant re-expression

Primary neonatal mouse cardiomyocytes were isolated from postnatal day 1-3 Phb2^flox/flox^ mice. Endogenous Phb2 was deleted by transduction with a Cre recombinase-expressing adenovirus (Ad-Cre, custom-built on the pAdEasy backbone, MOI 50). After 48 h, cells were transduced with recombinant adenoviruses expressing wild-type PHB2, PHB2-K253A/R255A or PHB2-I254G under identical conditions and at matched multiplicity of infection. Cells transduced with Cre alone served as the PHB2-deficient group. Loss of endogenous PHB2 and re-expression of the variant proteins at comparable levels were confirmed by immunoblotting before functional experiments. K253A/R255A and I254G were designed as helix-stabilizing and helix-destabilizing variants, respectively. Because structural predictions did not directly resolve the intended conformational states, the variants are interpreted throughout as perturbations of the PHB2 246-257 segment rather than as structurally validated helix-locked or coil-mimetic proteins.

### AlphaFold3 variant screening

Wild-type full-length mouse PHB2 (UniProt O35129; 299 amino acids) and seven target-segment variants were modelled as single-chain monomers using AlphaFold 3 v3.0.2. Mouse PHB2 was used because all cellular and in vivo experiments were performed in mouse cells and mice. Mouse and human PHB2 are both 299 residues, are identical across residues 236-266, and require no alignment offset, so residue numbering is interchangeable with the human numbering (UniProt Q99623) used for the structural analysis and the screen. The variants comprised K253A/R255A, K253Q/R255Q, K253E/R255E, A256G/A257G, I254G, A256P and R252P. The standard AlphaFold 3 data pipeline was used to generate protein multiple-sequence alignments and identify structural templates. Each construct was modelled using seeds 101, 202 and 303, with five diffusion samples generated per seed. The model with the highest ranking score within each seed was designated the seed-best model, yielding three primary models per construct and 24 primary models in total. All 120 generated models were retained for secondary analyses.

Structural analyses focused on residues 236-266, including the target segment spanning residues 246-257. Per-residue pLDDT values were obtained from the B-factor field of the predicted structure files, and regional pLDDT was calculated as the mean across the corresponding residues. Intra-segment predicted aligned error was calculated as the mean of the pairwise PAE values within residues 246-257. Secondary structure was assigned using the P-SEA implementation in Biotite^24^, and helix content was expressed as the fraction of target-segment residues assigned as α-helical.

Backbone hydrogen-bond compatibility was evaluated for all possible O(i)-N(i+4) pairs within each region. A pair was classified as geometrically compatible when the O-N distance was ≤3.5 Å. Pairs in which residue i+4 was proline were classified as chemically ineligible and scored as incompatible in the effective hydrogen-bond fraction.

Local bending was quantified as the angle between the principal axes obtained by singular-value decomposition of the Cα coordinates of residues 236-245 and 258-266. For each seed-best variant model, residues 236-245 and 258-266 were superposed onto the corresponding seed-best wild-type model using a Cα-based Kabsch fit. Target-segment deformation was subsequently quantified as the Cα RMSD of residues 246-257. Residue-level pLDDT and helix-assignment frequencies were calculated across the three seed-best models. Variant-level metrics were summarized using the median and interquartile range across these models.

Each variant was assigned one of three verdicts, shown in Extended Data Fig. 6a. The verdicts summarize six metrics reported in Extended Data Fig. 6b across the three seed-best models: target-segment pLDDT, intra-segment predicted aligned error, P-SEA helix fraction, effective i→i+4 hydrogen-bond fraction, flank-axis bend angle, and Cα RMSD relative to wild type. They were assigned by inspection of these metrics and were not defined by numerical thresholds; the three categories are defined in the legend of Extended Data Fig. 6.

AlphaFold 3 confidence metrics and derived geometric measurements were interpreted as model-based descriptors of the predicted folded state. They were not considered direct evidence of helix-coil energetics, conformational populations or temperature-dependent structural transitions.

### Immunofluorescence imaging and colocalization analysis

Cells were fixed with 4% paraformaldehyde for 15 min, permeabilized with 0.1% Triton X-100 for 10 min and blocked with 5% bovine serum albumin for 1 h at room temperature. Cells were incubated overnight at 4 °C with antibodies against TOM20 together with GABARAPL2, LC3A, LC3B, GABARAPL0 or GABARAPL1 (Supplementary Table 4), followed by species-matched Alexa Fluor-conjugated secondary antibodies. Nuclei were counterstained with DAPI. Images were acquired on a laser-scanning confocal microscope with a 63x oil-immersion objective. Z-stacks were collected at 0.3-µm intervals with identical laser power, detector gain, pinhole and pixel-resolution settings across all conditions.

Mitochondria-associated ATG8 signal was quantified in Fiji using the Coloc 2 plugin^25^ after background subtraction. Channel thresholds were set by the Costes automatic thresholding procedure rather than manually^26^. Manders’ M1^27^ was calculated as the fraction of ATG8-positive signal overlapping TOM20-positive mitochondria. Twenty cells per condition were analysed in each of six independent cardiomyocyte isolations, and cell-level values were averaged within each isolation so that the isolation, not the individual cell, was the unit of analysis (n = 6). Scale bars represent 10 µm. **Immunoblotting, co-immunoprecipitation and blue-native PAGE**

Whole-cell or cardiac-tissue lysates were analysed by SDS-PAGE and immunoblotting for the proteins indicated in the figures, including PHB2, PHB1, GABARAPL2, β-actin, VDAC1, LONP1, SOD2, MFN2, OPA1, phospho-DRP1 Ser616, DRP1, phospho-eIF2α Ser51, eIF2α, ATF4, CHOP, phospho-AMPK and AMPK. Antibodies, including host species, clonality, catalogue numbers and dilutions, are listed in Supplementary Table 4.

For co-immunoprecipitation, cells or cardiac tissues were lysed in ice-cold non-denaturing buffer containing 20 mM Tris-HCl (pH 7.4), 150 mM NaCl, 1 mM EDTA and 1% NP-40, supplemented with protease and phosphatase inhibitors. Clarified lysates were incubated with an anti-GABARAPL2 antibody (Supplementary Table 4) or control IgG overnight at 4 °C, followed by protein A/G agarose beads for 2 h. Beads were washed five times with lysis buffer, and bound proteins were eluted in SDS sample buffer and immunoblotted for PHB2.

Densitometry was performed in Fiji on unsaturated exposures, with local background subtracted using an adjacent region of the same dimensions as the measured band. For every co-immunoprecipitation, recovery was expressed as the double ratio:

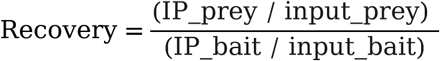

so that the recovered prey is normalized both to its own input and to the amount of bait captured in the same lane. Values were compared only within a single membrane; no comparison was made across blots. The blots in Fig. 3a and Fig. 4a are representative of three independent experiments; these panels are shown as representative blots and densitometry is not reported for them.

To assess PHB1-PHB2 complex formation, mitochondria-enriched fractions were solubilized with digitonin at a detergent-to-protein ratio of 4:1 (w/w) for 15 min on ice. After centrifugation at 20,000g for 30 min at 4 °C, supernatants were mixed with Coomassie G-250 sample additive, and 20 µg of protein was separated on 3-12% NativePAGE Bis-Tris gels at 150 V^28^. Dark-blue cathode buffer was replaced with light-blue cathode buffer after the dye front had migrated through one-third of the gel. Proteins were transferred to PVDF membranes at 25 V for 1 h and detected with antibodies against PHB1 and PHB2.

### Structure-aware mitochondrial LIR screen

A UniProt query based on the mitochondrial subcellular-location terms SL-0173 and SL-0172, retrieved 22 August 2026, returned 1,368 proteins^29^. Proteins with explicit mitochondrial inner- or outer-membrane annotations were retained, yielding 501 proteins. Intersection with the helix-LIR metric dataset identified 394 candidate motifs in 164 proteins, of which 108 retained explicit mitochondrial membrane annotation on review. Candidate motifs overlapping annotated transmembrane helices were excluded. The remaining motifs were required to have a mean helix pLDDT of at least 70 and a surface-exposed residue fraction of at least 0.50. Of the 394 motifs, 223 were excluded: 146 for lack of a mitochondrial membrane annotation, 63 for a surface-exposed fraction below 0.50 and 14 for a mean helix pLDDT below 70. This left 171 motifs in 95 proteins. One representative motif was retained per protein, producing a structural set of 95 protein representatives.

Because the two thresholds are arbitrary, the screen was repeated across all nine combinations of pLDDT (60, 70, 80) and surface exposure (0.40, 0.50, 0.60). PHB2 ranked 25th under every setting (Extended Data Fig. 2), so its position does not depend on the chosen cut-offs.

Structural eligibility was summarized by combining the surface-exposed residue fraction with the normalized mean helix pLDDT. This score reflects predicted structural accessibility and model confidence; it was not interpreted as folding stability, folding free energy or temperature-dependent unfolding. The iLIR database^11^ was used to cross-reference reported human LIR-containing proteins; PHB2 is indexed there under UniProt F5GY37 rather than under the canonical accession Q99623, and a query on Q99623 alone returns no record. The PHB2 YIKL motif is reported by iLIR with a PSSM score of 11 and a percentile of 40, where the percentile is the fraction of motifs scoring at or below that value. Nine of the 122 pages of the source table could not be retrieved because of server rate limiting; 5,600 of approximately 6,100 records (91.8%) were obtained. Residue positions were defined on the canonical UniProt PHB2 sequence Q99623. The analysed PHB2 motifs were YQRL at residues 121-124 and YIKL at residues 248-251.

### Structural analysis and geometric compatibility model

PHB2 residues 246-257 were examined in PHB1-PHB2 cryo-electron microscopy structures PDB 9O9Z, 9OA0 and 9UNL. DSSP assigned Gly247-Ala257 as α-helical in the displayed 9O9Z chain, with Pro246 in coil^30^. Cα distances between Tyr248 and Leu251 were measured across all 33 modelled PHB2 chains in the three structures. The 11 chains within each structure are related by the C11 symmetry of the prohibitin ring and are therefore not independent observations; the 33 values are reported as a description of the modelled ring, not as a sample of 33 independent measurements.

For the geometric-compatibility illustration, a frame from the λ = 0.79 trajectory with a Tyr248-Leu251 Cα distance of 9.63 Å was rigid-body superposed onto the backbone of the p62 LIR bound to LC3B (PDB 2ZJD). GABARAPL2 from PDB 7LK3 was then placed by superposition of its core onto LC3B. Hydrophobic pockets HP1 and HP2 were defined by receptor residues within 5 Å of p62 Trp340 and Leu343. No conformational search, scoring or energy minimization was performed; the model is an illustration of geometric compatibility, not a docking prediction.

### Molecular-dynamics simulations of PHB2 helicity

Replica-scaling molecular-dynamics simulations were used to test whether the LIR-bearing helix of PHB2 (residues 246-257) is conformationally more labile than a control helix in the same protein. Two fragments were simulated rather than the full-length protein, to concentrate sampling on the segment of interest while retaining its local structural context. The target fragment spans residues 190-267 and contains the LIR-bearing helix at residues 246-257 (fragment residues 57-68); the control fragment spans residues 81-189 and contains the control helix at residues 136-147. Both fragments were modelled with charged N- and C-termini (NH_3_^+^ and COO^-^) and standard pH 7 protonation states, with Asp and Glu deprotonated and Lys and Arg protonated.

Simulations were performed with GROMACS 2025.3^31^ and the CHARMM36m^32^ force field. Each fragment was solvated with TIP3P water in a rhombic-dodecahedral box. The net charge was neutralized with counter-ions, and the ionic strength was set to 150 mM KCl to approximate the potassium-rich intracellular environment of the mitochondrial intermembrane space and matrix, where PHB2 resides. The final systems contained 81,200 atoms (target) and 24,093 atoms (control). Each system was energy-minimized by steepest descent (5,000 steps) and equilibrated for 500 ps under NVT conditions at 340 K. Production runs were 200 ns in the NPT ensemble with a 2-fs time step. Covalent bonds to hydrogen were constrained with LINCS (fourth order, one iteration). van der Waals interactions used the force-switch modifier required by CHARMM36m, with switching from 1.0 nm, a cut-off at 1.2 nm and no dispersion correction. Long-range electrostatics were treated by particle-mesh Ewald summation (cubic interpolation, 0.12-nm grid). Temperature was maintained at 340 K with a velocity-rescaling thermostat (coupling constant 0.1 ps, protein and non-protein groups coupled separately), and pressure at 1 bar with an isotropic Parrinello-Rahman barostat (coupling constant 2.0 ps, compressibility 4.5 × 10^-5^ bar⁻¹). Coordinates were written every 10 ps, and initial velocities were drawn from a Maxwell distribution at 340 K with seed 1000.

A bath temperature of 340 K was used rather than 310 K in order to accelerate the sampling of unfolding events within the accessible trajectory length. This bath temperature alone did not destabilize either fragment: at λ = 1.0 the plateau helix fraction was 0.82 for the target and 0.76 for the control, so both helices remained folded under the unmodified force field. Because the target and control fragments were simulated under identical conditions, the comparison between them does not depend on the absolute bath temperature, which is not intended to represent a physiological or a heat-stress temperature.

Helical lability was probed by selectively scaling the force field of the helix of interest, which isolates the segment of interest from the rest of the fragment and avoids confounding helix-specific unfolding with global loss of structure. We applied the solute-tempering scheme of REST2, implemented with the partial_tempering routine of PLUMED 2.9.0^33^, without replica exchange. The scaling factor λ acts on the dihedral, Lennard-Jones and Coulomb terms of the target residues only: λ = 1.0 recovers the unmodified force field, and decreasing λ progressively destabilizes the helix. The target fragment was simulated at λ = 1.0, 0.93, 0.86, 0.79, 0.72, 0.65, 0.57 and 0.50, and the control fragment at λ = 1.0, 0.79 and 0.57. Each λ value was run as a single independent constant-λ trajectory, with no exchanges between λ values, so no exchange frequency or acceptance rate applies. In this scheme λ formally corresponds to an effective temperature of the scaled solute; the ladder was chosen to accelerate sampling of the helix-to-coil transition, and no numerical mapping from λ to a physical temperature is used anywhere in this study.

Because each λ is represented by a single trajectory, and because unfolding is effectively irreversible within the 200-ns window, these simulations establish relative lability and the accessibility of the required geometry; they do not yield a rate, a barrier or a temperature. The unfolding times annotated in Extended Data Fig. 3 are the last time at which the smoothed helix fraction exceeded 0.5 in each trajectory. They are descriptive values from single trajectories, not first-passage times or kinetic estimates, and they are not ordered by λ. Secondary structure was assigned with DSSP through MDTraj 1.10.3^34^. The helix fraction of the target (residues 246-257) or control (residues 136-147) segment was computed as the fraction of its residues assigned α-helical. Per-residue helix occupancy was used to localize the initiation of unfolding and was analysed over the first 30 ns to resolve the N-terminal Pro246-Gly247 initiation site. Backbone hydrogen-bond occupancy was calculated with the Baker-Hubbard criterion^35^, and salt bridges were defined by a nitrogen-oxygen distance below 4 Å between Lys/Arg and Asp/Glu side chains. Solvent-accessible surface area was computed with the Shrake-Rupley algorithm^36^. To assess access to the extended, binding-competent geometry, the Cα distance between the two LIR anchors Tyr248 and Leu251 was measured over each trajectory and compared with the 8.99 ± 1 Å separation observed in ATG8-LIR complexes. Analyses treated individual frames as within-trajectory descriptions and did not assume convergence of the folded ensemble at reduced λ.

### Molecular-dynamics simulations of PHB2-ATG8 complexes

Complexes comprising the PHB2-derived peptide NPGYIKLRK and LC3A, LC3B, GABARAPL0, GABARAPL1 or GABARAPL2 were prepared using tleap in AmberTools 26^37,38^. The peptide corresponds to PHB2 residues 245-253 and contains the YIKL motif at residues 248-251; it is a nine-residue subset of the 246-257 segment used in the fragment simulations, chosen to span the LIR core and its immediate flanks in the geometry adopted on binding. Protein and peptide atoms were described using the Amber ff14SB force field^39^. Each complex was solvated in a truncated-octahedral box of TIP3P water extending at least 10 Å from the solute, neutralized with Na^+^ or Cl^-^ ions and supplemented with approximately 0.15 M NaCl using ion parameters compatible with TIP3P water. The resulting systems contained 19,285-24,046 atoms and 5,718-7,294 water molecules. Before simulation, each topology was verified to contain the complete nine-residue NPGYIKLRK peptide and to pass atom-count, residue-count, close-contact and GPU smoke-test checks.

All simulations were performed using the GPU-accelerated pmemd.cuda implementation in Amber 26. Energy minimization comprised 5,000 cycles with a 10 kcal mol^-1^ Å^-2^ positional restraint on the receptor-peptide complex, followed by 10,000 cycles of unrestrained minimization; the first half of each minimization used steepest descent and the second half used conjugate gradients. Systems were heated from 0 to 300 K over 0.5 ns under constant-volume conditions using a Langevin thermostat with a collision frequency of 2 ps^-1^ and a 5 kcal mol^-1^ Å^-2^ restraint on the solute. Density equilibration was then performed at 300 K and 1 bar, beginning with five consecutive 20-ps Berendsen-barostat blocks with a 2 kcal mol^-1^ Å^-2^ solute restraint, followed by fifty 20-ps unrestrained blocks using the Monte Carlo barostat. This was followed by 1 ns and 3 ns of unrestrained NPT equilibration using the Monte Carlo barostat, giving a total pre-production simulation time of 5.6 ns.

Production simulations were performed in the NPT ensemble at 300 K and 1 bar using the Langevin thermostat and Monte Carlo barostat, with thermostat and pressure-relaxation constants of 2 ps^-1^ and 2 ps, respectively. These simulations were run at 300 K, the standard reference temperature for the ff14SB parameter set, because they were used to compare the relative stability of the five receptor complexes under identical conditions rather than to model a physiological or a heat-stress temperature. A 2-fs integration time step was used, and bonds involving hydrogen atoms were constrained using SHAKE. Long-range electrostatic interactions were treated using particle-mesh Ewald summation, with a 10-Å real-space non-bonded cut-off. Centre-of-mass motion was removed every 1,000 steps. Each 200-ns trajectory was divided into twenty sequential 10-ns segments to permit checkpointed continuation. Coordinates were written in NetCDF format every 20 ps, thermodynamic quantities every 100 ps and restart files every 1 ns. Independent simulations used distinct Langevin random-number seeds.

For trajectory analysis, the first 200 ns of two independent trajectories for each receptor were sampled at 1-ns intervals, yielding 200 frames per replicate. Periodic-boundary artefacts were corrected by unwrapping the receptor and peptide separately and translating the peptide to its nearest periodic image relative to the receptor. Trajectories were aligned to their initial frames using receptor Cα atoms. Receptor and peptide RMSD, radius of gyration, receptor-peptide centroid and minimum heavy-atom distances, interface contacts, native-contact fraction and per-residue RMSF were calculated. Contacts were defined by a 4.5-Å heavy-atom cut-off. Receptor-aligned peptide conformations were further examined using principal-component analysis and average-linkage hierarchical clustering with a 2.0-Å threshold. Independent trajectories were treated as the statistical unit (n = 2); individual frames were used only for within-trajectory characterization. With two trajectories per receptor, between-receptor differences are described rather than tested. Trajectory processing and analysis were performed using Python and MDAnalysis^40^.

### Recombinant-protein expression and temperature-dependent binding

Flag-MBP-tagged LC3A, LC3B, GABARAPL0, GABARAPL1 and GABARAPL2, and His-MBP-tagged PHB2 residues 190-299, were expressed in *E. coli*. Flag-tagged receptors were captured on anti-FLAG M2 affinity resin (Sigma-Aldrich) and eluted by competition with 3xFLAG peptide (Beyotime); His-MBP-PHB2(190-299) was purified on Ni-NTA resin (GE Healthcare). All purification was performed at 4 °C in 20 mM HEPES pH 7.5, 100 mM NaCl, 10% (v/v) glycerol and 0.00225% (w/v) lauryl maltose neopentyl glycol (LMNG), and proteins were used immediately after purification without storage or freeze-thaw. Purified proteins were not subjected to size-exclusion chromatography, and their oligomeric state and monodispersity were not assessed independently.

For temperature-dependent co-immunoprecipitation, equimolar amounts of receptor and PHB2 fragment, at 10 uM each, were incubated for 30 min at 293, 298, 303, 308, 313 or 318 K. Specificity controls omitting the receptor or the PHB2 fragment were run at 293 K and, in the reversibility experiment described below, at 318 K, so that non-specific capture was measured at both ends of the temperature range. Complexes were recovered through the Flag-tagged ATG8 protein and immunoblotted for both partners. Input samples and recovered-bait blots were collected at every temperature to confirm that both proteins were present at comparable levels and that equivalent amounts of bait were captured across the series (Extended Data Fig. 5).

Because the receptors were eluted by peptide competition and were not buffer-exchanged afterwards, free 3xFLAG peptide was carried into the binding reaction and the subsequent capture step. The peptide was present at the same concentration in every sample of a temperature series, and recovery was normalized to the amount of bait actually captured in each lane, so any reduction in capture efficiency caused by the free peptide applies equally across the series and is divided out by that normalization. Recovered-bait blots collected at every temperature (Extended Data Fig. 5) confirm that comparable amounts of bait were captured across the series.

Recovered PHB2 was quantified by densitometry as described above, as the double ratio (IP_PHB2/input_PHB2)/(IP_bait/input_bait), with local background subtracted from each band using an adjacent region of the same dimensions. Each temperature series was performed in three independent experiments.

The thermal stability of the individual proteins was not assessed separately across the temperature series, so a contribution of temperature-dependent aggregation or degradation of either partner to the measured recovery cannot be excluded directly. The reversibility experiment described below bears on this: recovery at 313 K returned to the 293 K level when the sample was returned to 293 K, which is not expected if the increase at 313 K reflected irreversible aggregation of either protein.

Reversibility was tested by forming complexes at 313 K or 318 K, returning them to 293 K and re-assaying binding, with samples held continuously at 293 K, 313 K or 318 K assayed in parallel on the same membrane. A no-receptor control was included at 318 K. Recovery was quantified by the same double ratio and analysed in three independent experiments, with five preselected comparisons corrected by the Holm-Šídák method^41^.

### Mitochondrial-respiration measurements

Cellular oxygen-consumption rate (OCR) was measured with a Seahorse XFe96 extracellular-flux analyser (Agilent). Cardiomyocytes were seeded at 2.0 × 10^4^ cells per well in XFe96 microplates and assayed 48 h later in XF base medium supplemented with 10 mM glucose, 1 mM pyruvate and 2 mM glutamine (pH 7.4), after a 1-h incubation at 37 °C without CO_2_. Oligomycin (1.5 µM), FCCP (1.0 µM) and rotenone plus antimycin A (0.5 µM each) were injected sequentially following the manufacturer’s Mito Stress Test protocol. Basal respiration was calculated as baseline OCR minus non-mitochondrial OCR, and maximal respiration as peak FCCP-stimulated OCR minus non-mitochondrial OCR. All OCR values were normalized to total protein content, determined by BCA assay (Pierce) in the same wells immediately after each run. Technical replicate wells were averaged within each independent cardiomyocyte isolation, so that the isolation was the unit of analysis; eight independent isolations were assayed per condition (n = 8). Data are mean ± s.e.m. Because every isolation contributed to both constructs and both temperatures, differences were assessed by repeated-measures two-way ANOVA with matching on both factors, with the construct × temperature interaction as the primary test, followed by Tukey’s multiple-comparison test across the group means.

### PHB2 K253A/R255A mouse model and experimental heat stress

Cardiomyocyte-specific PHB2 K253A/R255A knock-in mice were generated using a Cre-dependent conditional knock-in strategy. A targeting cassette containing a loxP-flanked wild-type mouse Phb2 coding sequence followed by a 3xSV40 polyadenylation signal was inserted immediately upstream of the endogenous Phb2 start codon, and the K253A and R255A substitutions were introduced into the downstream endogenous coding sequence. Cas9 mRNA, guide RNA and the targeting vector were co-injected into fertilized C57BL/6N eggs. Founder mice were identified by PCR genotyping and confirmed by Sanger sequencing. Homozygous conditional knock-in mice were crossed with Myh6-Cre transgenic mice^42^. In cardiomyocytes, Cre-mediated excision of the floxed wild-type cassette activates expression of the endogenous PHB2 K253A/R255A allele. Myh6-Cre+; Phb2^K253A/R255A-fl/fl^ mice were used as the mutant group, and Cre-negative Phb2^K253A/R255A-fl/fl^ littermates served as controls. Control littermates express wild-type PHB2 from the unexcised cassette and are referred to as wild-type throughout for readability.

Male and female mice aged 6-8 weeks were used, with four males and three females in each group. In the mutant heat-stroke group, the animal that died was male, leaving three males and three females in that group at the time of sampling. Group sizes were not designed to detect differences between sexes, and sex was not included as a factor in the analyses. Mice were housed at 295 ± 1 K (22 °C) and 65-70% relative humidity under a 12-h light-dark cycle, with free access to food and water. Genotype was determined by PCR before allocation. After acclimation, mice of each genotype were allocated to the heat-stress or the sham group by a computer-generated random-number sequence, stratified by sex so that each group contained four males and three females. Allocation to treatment was therefore randomized within genotype and within sex. Genotype itself is determined by breeding and was not randomized. Sample size was set at seven mice per group on the basis of group sizes used in previous murine heat-stroke studies and was not determined by a formal power calculation. Effect sizes are reported with 95% confidence intervals throughout.

Heat-stress animals were placed in a temperature-controlled chamber at 314.4 ± 0.2 K and 50 ± 5% relative humidity for a fixed exposure of 58 min, after which they were returned to room temperature for recovery. Core body temperature was recorded with a rectal probe every 2 min throughout the exposure and reached approximately 315 K by the end of the exposure in all heat-stressed animals. Sham-treated mice underwent identical handling at 295 ± 1 K without hyperthermia. Blood was collected before heat exposure (0 h) and at 3, 6 and 24 h after recovery. Echocardiography was performed at 24 h, after which mice were euthanized and cardiac tissue collected for biochemical and metabolomic analyses. All animal procedures were approved by the Animal Ethics Committee of the Second Affiliated Hospital, Zhejiang University School of Medicine (approval number 2025-281), and were performed in accordance with institutional guidelines for the care and use of laboratory animals.

One mouse in the mutant heat-stroke group died after the procedure before sampling. Because this animal was plausibly the most severely affected, its death was treated as potentially informative missingness: it is reported in the figure legend and the Results, group sizes are n = 6 (mutant heat stroke) and n = 7 (all other groups), and the direction of the resulting bias, towards the null, is stated where the affected endpoints are described. No other animals or data points were excluded.

### Echocardiography and circulating-injury markers

Cardiac function was assessed with a Vevo F2 ultrasound imaging system (FUJIFILM VisualSonics) fitted with a UHF57x transducer (25-57 MHz). Mice were anaesthetized with isoflurane, 3% for induction and 1.5% for maintenance, and positioned on a heated platform to maintain body temperature. Imaging began only once the heart rate had stabilized. Parasternal long-axis and short-axis views were acquired in B-mode and M-mode. Heart rate, left-ventricular internal dimension at end-diastole and end-systole, anterior and posterior wall thicknesses, ejection fraction and fractional shortening were measured, and left-ventricular mass was computed from these measurements by the Vevo LAB software. At least three consecutive cardiac cycles were measured per animal in Vevo LAB. Heart rate was recorded for every animal. Images were acquired and measured by the same operator, who was unaware of both the genotype and the treatment assignment of the animals throughout acquisition and analysis. Cardiac troponin I was measured at 0, 3, 6 and 24 h after heat exposure with a mouse TNNI3/cTn-I enzyme-linked immunosorbent assay (Elabscience, E-EL-M1203). Serum was diluted 1:5 before assay; the kit has a stated standard range of 7.81-500 pg mL^-1^, giving an effective range of 0.039-2.5 ng mL^-1^ for the diluted samples. The lower limit of quantification was set at 0.05 ng mL⁻¹; values below it were assigned that limit, and the number of such values in each group is stated in the figure legend. Serum alanine aminotransferase, aspartate aminotransferase, creatinine and urea nitrogen were measured on an IDEXX Catalyst One veterinary chemistry analyser using the manufacturer’s dry-slide reagents. These are enzymatic and colorimetric assays rather than immunoassays, and the analyser is intended for animal samples. Serum IL-6, TNF-α and IL-1β were measured with mouse ELISA kits (Thermo Fisher Scientific; KMC0061, BMS607-3 and BMS6002, respectively). These seven markers were analysed as a preselected family and P values were corrected across them by the Holm method.

### Cardiac metabolomics

Cardiac tissue from all 27 mice in the study was subjected to targeted metabolomic profiling by Cosmos Wisdom Biotech (Hangzhou, China) using the Q300 quantitative targeted metabolomics kit, quantifying 184 metabolites, with analysis focused on 19 carnitine species (free carnitine and 18 acylcarnitines). Tissue was homogenized with zirconia beads and extracted with ice-cold methanol containing stable-isotope-labelled internal standards according to the manufacturer’s protocol, and metabolite concentrations were normalized to tissue wet weight. Extracts were derivatized with freshly prepared 3-nitrophenylhydrazine and 1-ethyl-3-(3-dimethylaminopropyl)carbodiimide for 60 min at 30 °C to detect carboxyl-containing metabolites, diluted with ice-cold 50% methanol, held at -20 °C for 20 min and centrifuged. Supernatants were transferred to a 96-well plate, spiked with isotope-labelled internal standards, and analysed on a Waters ACQUITY UPLC system coupled to a Xevo TQ-S triple-quadrupole mass spectrometer. Data were processed with MassLynx v4.1. Group sizes were n = 7 (wild-type sham), 7 (mutant sham), 7 (wild-type heat stroke) and 6 (mutant heat stroke), the smaller final group reflecting the single death described above. Sample identifiers in the source workbook are mouse ear tags and do not encode group assignment; group membership was taken from a separate annotation file and verified before analysis. No sample was excluded.

Metabolite intensities were log2-transformed, with zero values replaced by half the smallest positive value for that metabolite. Each metabolite was then standardized to a z-score across all 27 samples, so that the standardization set and the analysis set are identical, and the aggregate acylcarnitine score was defined as the mean z-score across the 19 carnitine species. Between-group differences (mutant heat-stressed versus wild-type heat-stressed) are expressed as log2 fold changes. The acylcarnitine score was analysed by two-way ANOVA with the genotype × heat-stroke interaction as the primary test. Individual carnitine species were compared by two-tailed Welch’s t-test; the species highlighted in Fig. 4i are those reaching uncorrected P < 0.05 and are reported as exploratory, hypothesis-generating comparisons rather than as findings that survive correction across the 19 species. Benjamini-Hochberg false-discovery-rate correction was also computed across all 184 metabolites.

### Statistical analysis

Data are presented as mean ± s.e.m. unless otherwise stated, with individual data points shown wherever n permits. For experiments in which the same independent cardiomyocyte isolations were measured under both conditions (colocalization and respiration), data were analysed by repeated-measures two-way ANOVA with matching on both factors, with the interaction term (temperature × bafilomycin, or construct × temperature) as the primary test; pairwise comparisons of cell means were made with a common error term and Šídák’s or Tukey’s multiple-comparison correction, as indicated in the figure legends. Fold changes of ATG8-family paralogues over their matched 310 K controls were tested on a log scale against unity and corrected across the five paralogues by the Holm method. In the reversibility experiment, five preselected comparisons were corrected by the Holm-Šídák method. Animal endpoints were analysed by two-way ANOVA, with the genotype × heat-stroke interaction as the primary test, followed by Tukey’s test across the four group means. Serum cardiac troponin I was analysed with a mixed-effects model with matching by time; because half of the sham values were at the lower limit of quantification, statistical comparison was restricted to the two heat-stroke groups, with Šídák correction across the four time points. The seven serum injury and inflammation markers were corrected by the Holm method across that family. Effect sizes are reported with 95% confidence intervals, and exact P values are reported where possible; P < 0.05 was considered statistically significant. For cell experiments, independent cardiomyocyte isolations were treated as biological replicates; for animal experiments, each mouse represented one biological replicate. Statistical tests were performed with GraphPad Prism 11.0.1 and with Python 3.9.6 using SciPy 1.13.1, NumPy 2.0.2, pandas 2.3.3 and statsmodels 0.14.6.

## Data availability

The structures used in this study are available from the Protein Data Bank under accessions 9O9Z, 9OA0 and 9UNL (PHB1-PHB2 complexes), 8Q6Q (GABARAPL2-IRGQ), 2ZJD (LC3B-p62) and 7LK3 (GABARAPL2). Sequences and predicted models were obtained from UniProt under accessions Q99623 (human PHB2), O35129 (mouse PHB2) and F5GY37, the entry under which PHB2 is indexed in the iLIR database. Source data for every main and Extended Data figure panel, including all densitometry values, per-well respirometry measurements, per-animal echocardiography and serum measurements, and per-sample metabolite intensities, will be provided with the final version of the manuscript. Uncropped images of all immunoblots shown in the figures are provided in Supplementary Fig. 1. Targeted metabolomics data will be deposited in MetaboLights before publication and the accession will be provided at that time. Molecular-dynamics input files, topologies, parameter and run-control files, the retrieved iLIR records, the structure-aware screen output and all AlphaFold 3 models will be deposited at Zenodo before publication. Production trajectories are large and will be made available from the corresponding author on request. All other data supporting the findings of this study are available within the paper and its Supplementary Information.

## Code availability

Analysis and figure-generation code will be deposited at Zenodo before publication. This includes GROMACS and PLUMED run configurations, Amber input files, the MDTraj and MDAnalysis analysis scripts, the structure-aware screen pipeline, and the Python scripts that generate each figure panel. No custom algorithms were used beyond those described in the Methods. Code is available from the corresponding author on request in the interim.

---

**Extended Data Fig. 1.**
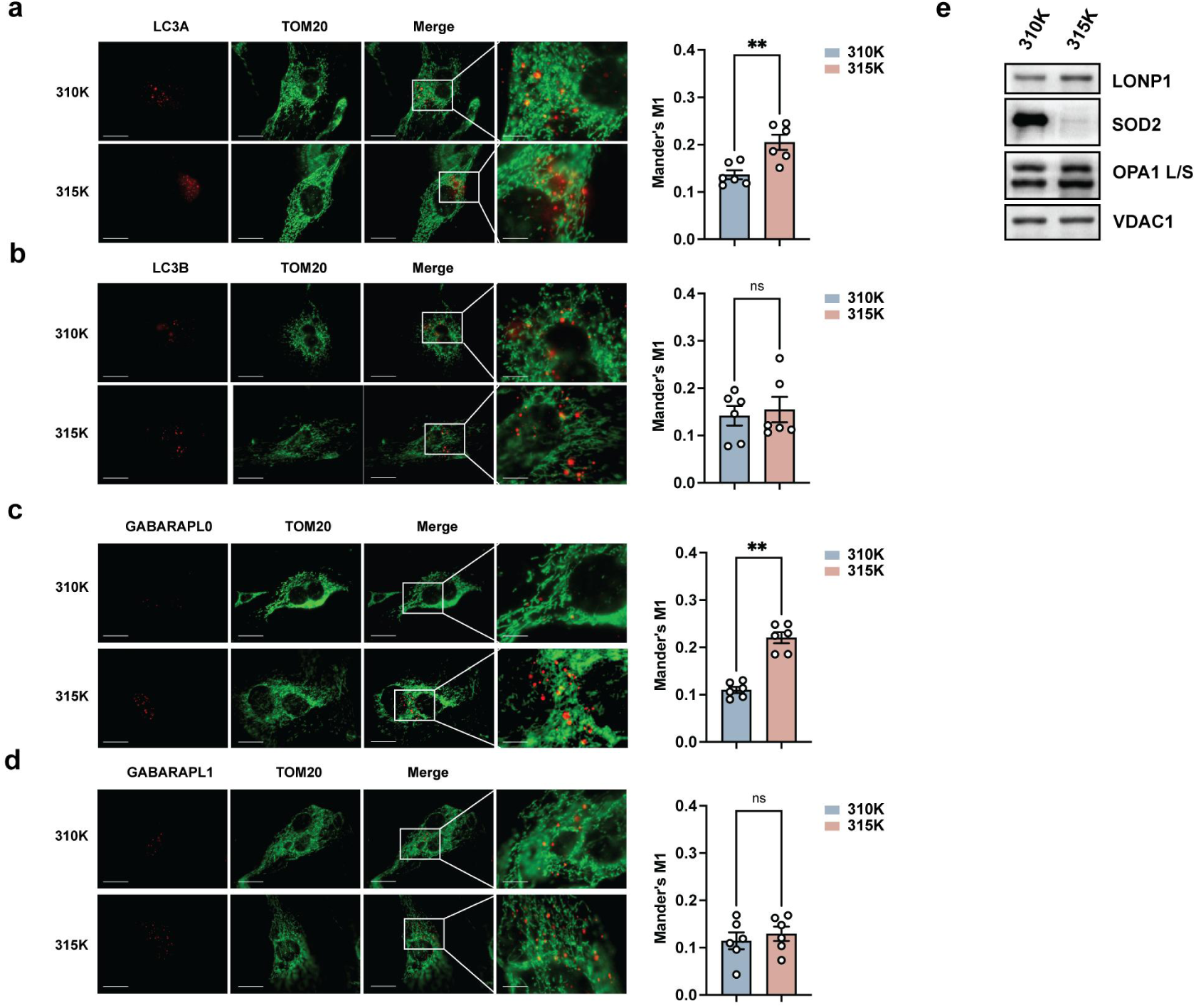
Colocalisation of the remaining four ATG8-family proteins, and mitochondrial stress markers. **a–d,** Immunofluorescence colocalisation of LC3A (**a**), LC3B (**b**), GABARAPL0 (**c**) and GABARAPL1 (**d**) (red) with TOM20 (green) in primary neonatal cardiomyocytes held at 310 K or 315 K. The fourth column shows the boxed region enlarged; the right-hand panel in each row is the quantification, as Mander’s M1 with Costes automatic thresholds as in Fig. 1b. Bars are mean ± s.e.m.; points are independent cardiomyocyte isolations (n = 6, 20 cells each), with statistics computed on isolation-level means. Fold changes over the matched 310 K control were tested on a log scale against unity and corrected across the five ATG8-family paralogues by the Holm method, as in Fig. 1c; the fifth paralogue, GABARAPL2, is shown in Fig. 1a,b. \*\**P* < 0.01; ns, not significant. Scale bars, 10 μm (2 μm in enlargements). e, Immunoblots of cardiomyocytes held at 310 K or 315 K, probed for LONP1, SOD2 and the long and short forms of OPA1, with VDAC1 as a mitochondrial loading control. Representative of n = 3 independent isolations.

**Extended Data Fig. 2.**
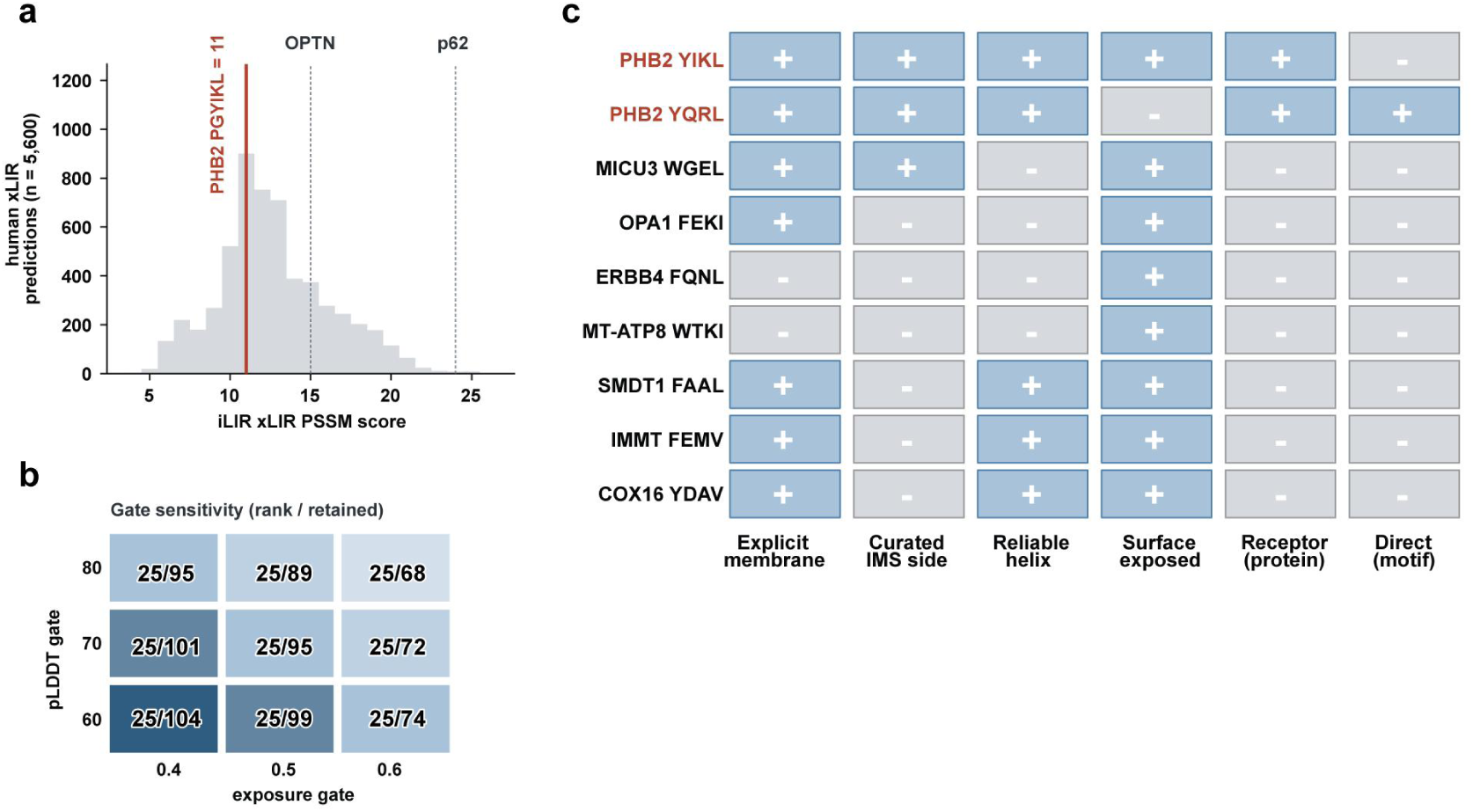
Motif provenance, gate sensitivity and the candidate evidence matrix. **a,** Distribution of iLIR xLIR PSSM scores across human predictions (iLIR Autophagy Database, Homo sapiens LIRCP table, retrieved 22 August 2026). The distribution is based on 5,600 of approximately 6,100 records; nine of the 122 source pages were unreachable because of server throttling. Position in the retrieved table is unrelated to motif score (Spearman ρ = -0.013 between record position and PSSM score, *P* = 0.32; per-page mean scores 10.9 to 16.0 about an overall mean of 12.7), so the missing pages are not expected to shift the distribution. The red line marks the PHB2 6-mer PGYIKL (residues 246–251, score 11); dotted lines mark two experimentally validated LIRs, OPTN DSFVEI (score 15) and p62/SQSTM1 DDWTHL (score 24), for reference. **b,** Sensitivity of the two structural gates across nine combinations of thresholds (helix pLDDT 60, 70 or 80; surface-exposed fraction 0.4, 0.5 or 0.6). Each cell gives the rank of PHB2 YIKL 248–251 over the number of candidate proteins retained at that setting; the combination used throughout is pLDDT ≥ 70 with exposure ≥ 0.5. **c,** Evidence matrix for nine representative candidate sites: four retained (PHB2 YIKL, SMDT1, IMMT and COX16) and five excluded, illustrating each exclusion criterion. Two of the excluded sites were prespecified checks, ERBB4 FQNL expected to fail on membrane annotation and OPA1 FEKI on helix confidence. Columns are, in order: explicit membrane annotation; curated intermembrane-space-side assignment; helix confidence above the pLDDT gate; surface exposure above the exposure gate; receptor evidence at the protein level; and direct evidence at the motif level. Plus signs indicate the criterion is met, minus signs that it is not. The two PHB2 sites are marked in red.

**Extended Data Fig. 3.**
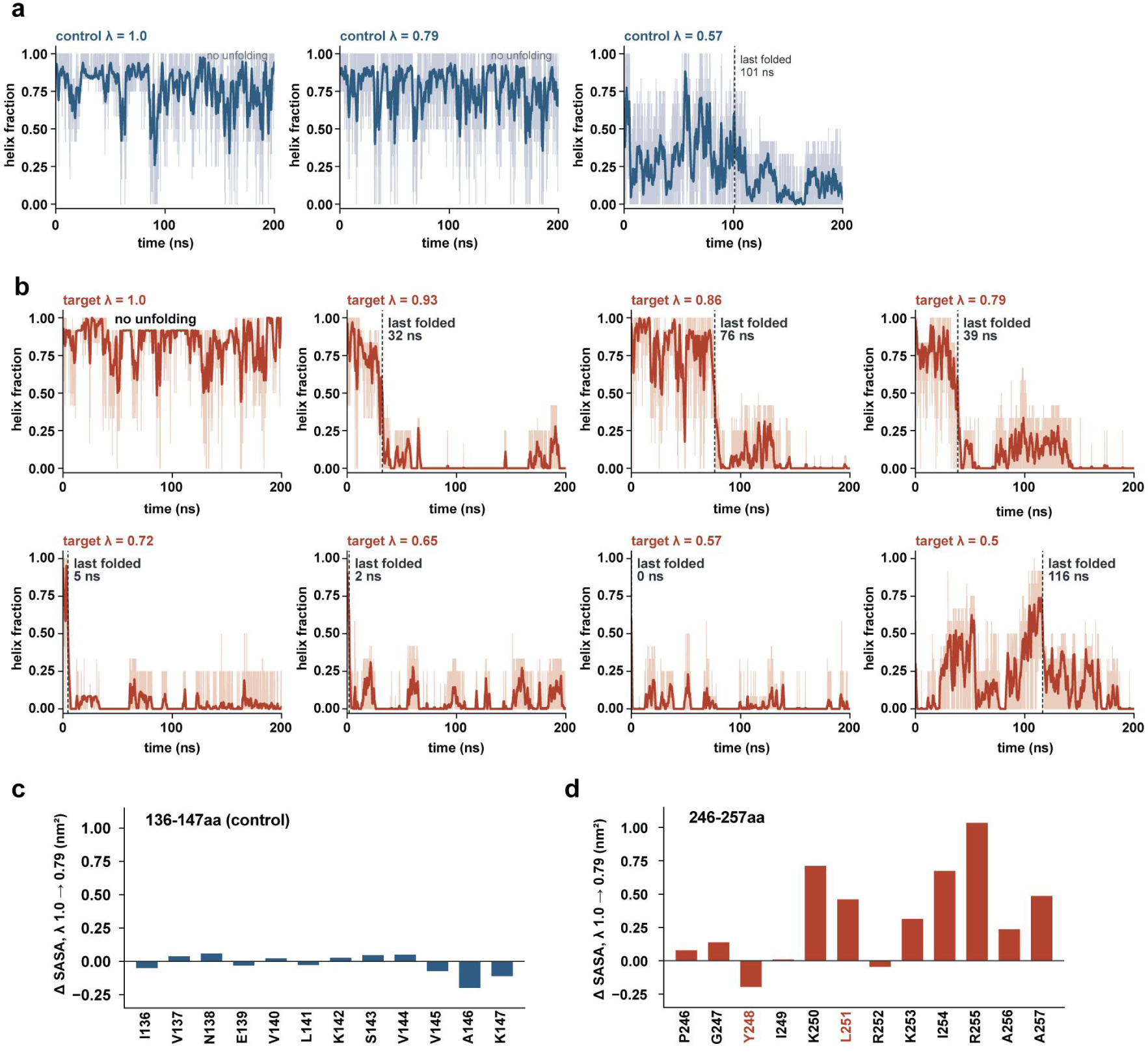
Unfolding of the target and control segments across the λ ladder, and the accompanying change in solvent exposure. **a,** Helical fraction of the control segment, PHB2 136–147, at λ = 1.0, 0.79 and 0.57. The pale trace is the per-frame value, the bold trace a 5 ns running mean, and the dashed line marks the last time the smoothed fraction exceeded 0.5. **b,** The same for the target segment, PHB2 246–257, at eight values of λ; the control was run at three, spanning the range over which the target unfolds. The annotated times use the same definition as in a and are taken from single trajectories, and are not ordered by λ; at λ = 0.5 the segment begins the trajectory largely unfolded and gains helicity after roughly 100 ns, so its annotation reflects late helix formation rather than prolonged resistance to unfolding. **c,d,** Change in per-residue solvent-accessible surface area (nm², λ = 0.79 minus λ = 1.0) for the control (**c**) and target (**d**) segments. The two LIR anchor residues, Tyr248 and Leu251, are marked in red in **d**.

**Extended Data Fig. 4.**
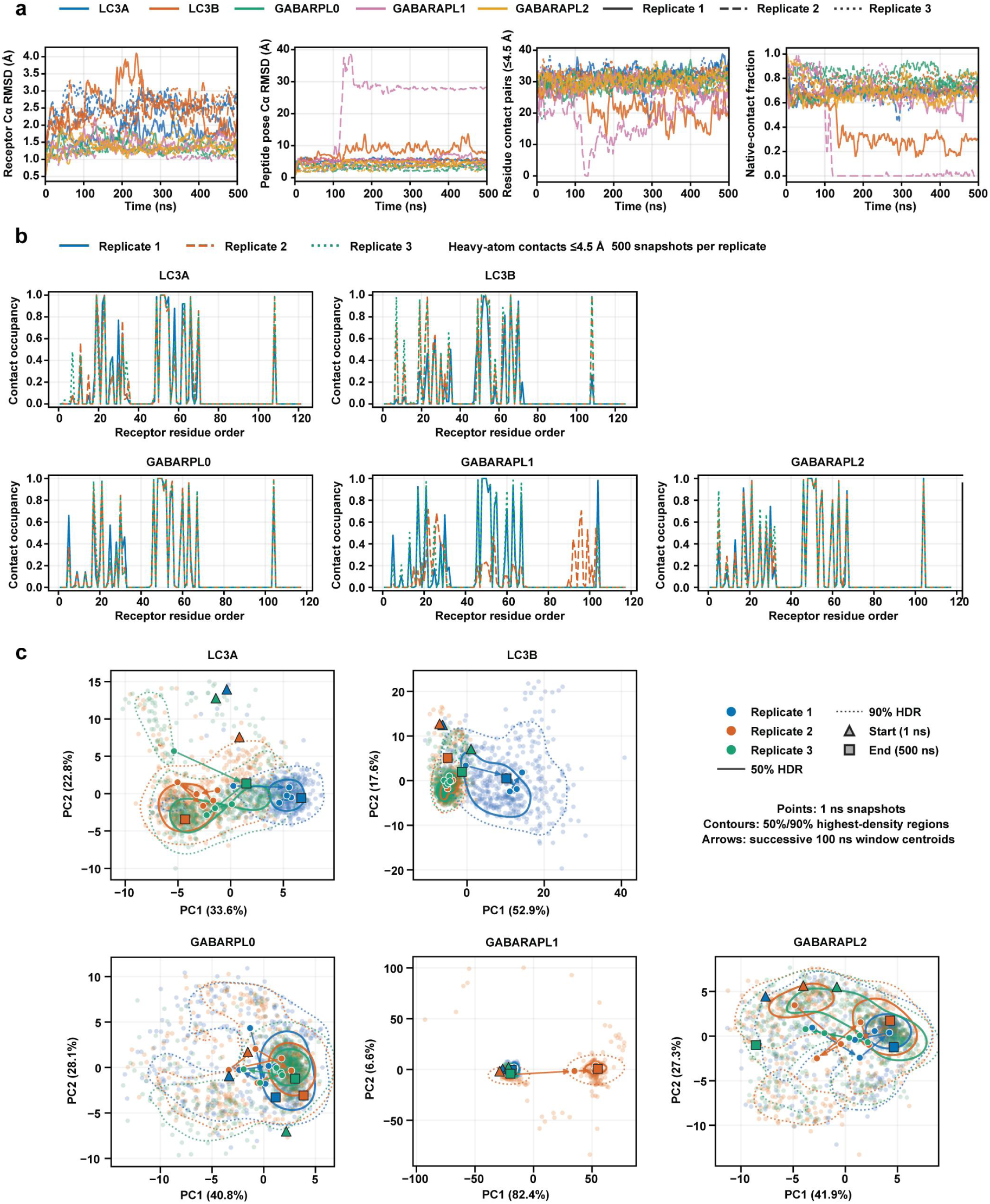
The PHB2 LIR peptide bound to five ATG8-family proteins. **a,** Trajectory stability for the PHB2-derived peptide NPGYIKLRK (residues 245-253) bound to LC3A, LC3B, GABARAPL0, GABARAPL1 and GABARAPL2, in three 500 ns simulations per system (15 trajectories; frames sampled every 1 ns). Left to right: receptor Cα RMSD; peptide pose Cα RMSD; number of receptor–peptide residue contact pairs (heavy-atom distance ≤ 4.5 Å); and native-contact fraction, referenced to the first frame of each replicate. Colour denotes receptor, line style denotes replicate. The peptide remains bound throughout in LC3A, GABARAPL0 and GABARAPL2. In GABARAPL1 replicate 2 it leaves the binding groove at 100–125 ns; that trajectory is truncated at 120 ns, because beyond this point the peptide–receptor centroid separation exceeds the minimum-image limit of the simulation box (32.8 Å for a 65.6 Å box) and the coordinates are no longer interpretable. **b,** Per-position contact occupancy at the receptor interface, computed over 500 snapshots per replicate using the same 4.5 Å heavy-atom criterion, with all three replicates overlaid (replicate 1, blue solid; replicate 2, orange dashed; replicate 3, green dotted). The x axis is a positional index along the receptor rather than residue numbering, so it is comparable within a panel but not between panels. **c,** Principal component analysis of the bound peptide pose, performed separately for each system, so the principal-component directions are not comparable between panels. Axis units are Å; percentages give the variance explained.

**Extended Data Fig. 5.**
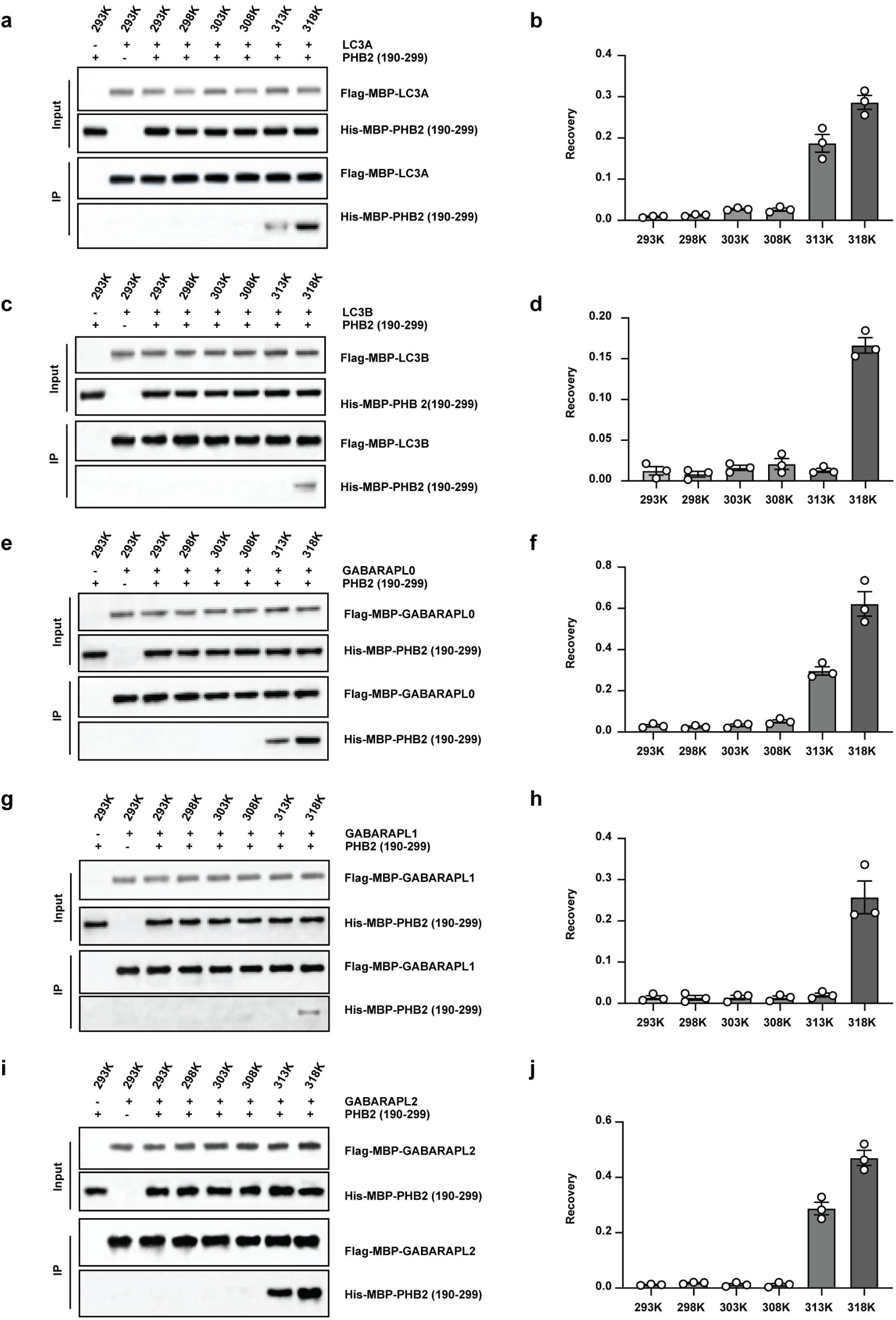
Temperature dependence of PHB2 binding for the five ATG8-family proteins. **a,c,e,g,i,** *In vitro* co-immunoprecipitation of Flag-MBP-tagged LC3A (**a**), LC3B (**c**), GABARAPL0 (**e**), GABARAPL1 (**g**) and GABARAPL2 (**i**) with His-MBP-PHB2(190–299) at 293–318 K. Each panel shows input and recovered Flag-MBP-receptor and His-MBP-PHB2; the recovered PHB2 row is the readout of binding, the others serving as loading and bait-capture controls. The first two lanes of each panel are specificity controls at 293 K, omitting the receptor and the PHB2 fragment respectively. **b,d,f,h,j,** Densitometric quantification of **a**, **c**, **e**, **g** and **i**. Recovery is the ratio of recovered to input PHB2, divided by the ratio of recovered to input receptor, so that it is corrected for both loading and bait capture. Background was subtracted using a region of the same size immediately adjacent to each band; the residual values at 293–308 K, where no band is visible, indicate the background level of the measurement. Bars are mean ± s.e.m.; points are independent experiments (n = 3). Each receptor was assayed and quantified on a separate blot, so recovery values are comparable between temperatures within a panel but not between panels; the comparison between receptors is made on the single blot in Fig. 2h,i.

**Extended Data Fig. 6.**
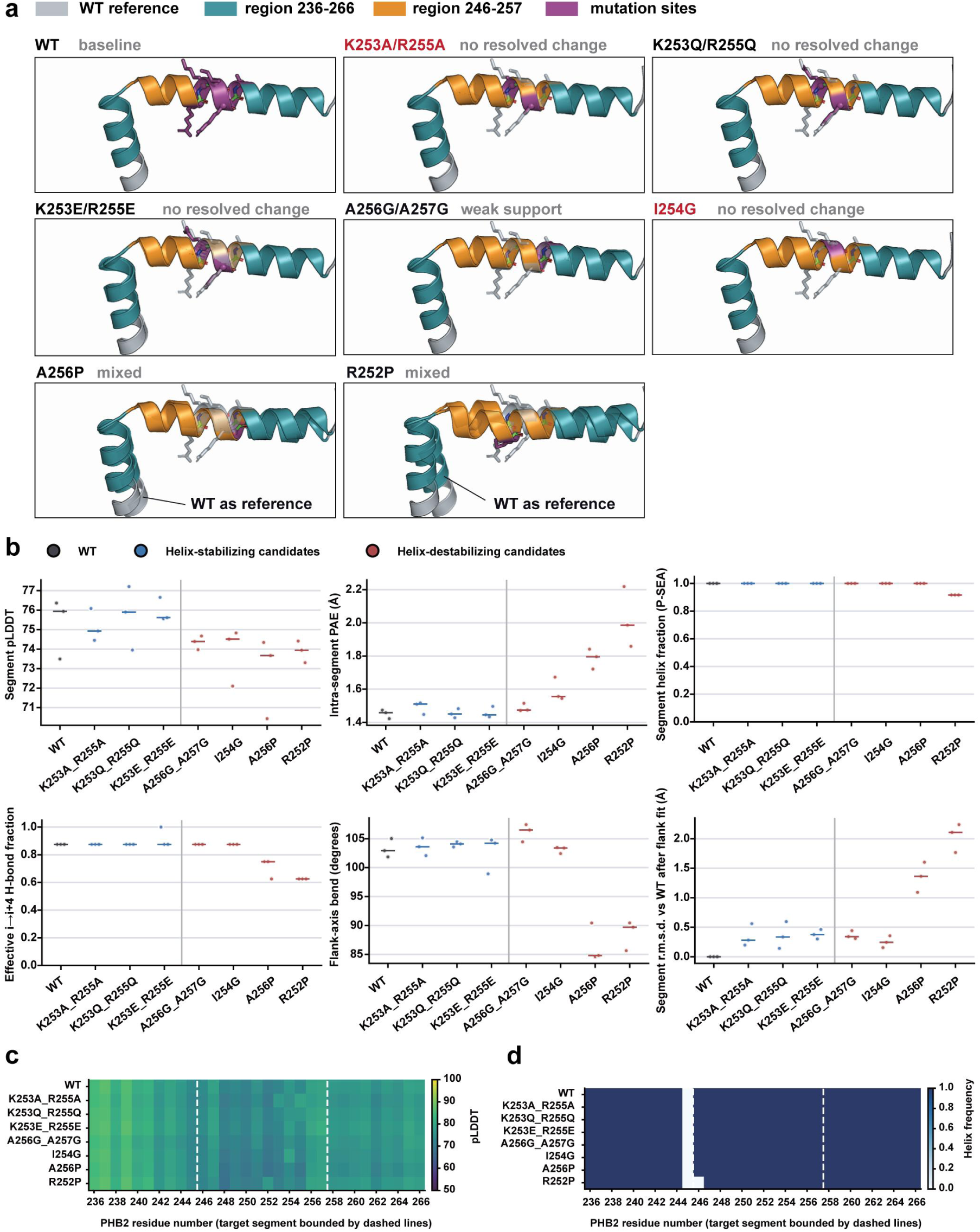
AlphaFold3 screen of candidate substitutions in the PHB2 target segment. Models are of mouse PHB2 (UniProt O35129), which is identical to human PHB2 across residues 236–266, so the numbering is the same as elsewhere in the paper. **a,** Predicted structures of wild-type PHB2 and seven candidate substitutions. Teal marks residues 236–266, orange the target segment 246–257, magenta the substituted positions, and grey the superposed wild-type reference. Each panel is annotated with the verdict for that substitution; the two taken forward are labelled in red. **b,** Six structural metrics for the target segment. Each point is the best model from one seed (three seeds per substitution) and the horizontal line is the mean. Top row, left to right: mean pLDDT of the segment; predicted aligned error within the segment; helix fraction of the segment assigned by P-SEA. Bottom row: effective i→i+4 hydrogen-bond fraction; bend angle between the axes of the two flanking helices; and r.m.s.d. of the segment relative to wild type after superposition on the flanking regions. Substitutions are grouped by intended effect, helix-stabilising (blue) or helix-destabilising (red), separated by the vertical line. The verdicts in a summarise these metrics: "no resolved change" where every metric overlaps the wild-type range across the three seeds; "weak support" where some metrics shift in the intended direction but not consistently across seeds or metrics; and "mixed" where metrics change but the change extends to the flanking geometry rather than being confined to the target segment. **c,** Per-residue pLDDT across residues 236–266, as the mean of the three seed-best models. Dashed lines bound the target segment. **d,** Frequency with which each residue is assigned as helical across the three seed-best models. Dashed lines bound the target segment.

**Extended Data Fig. 7.**
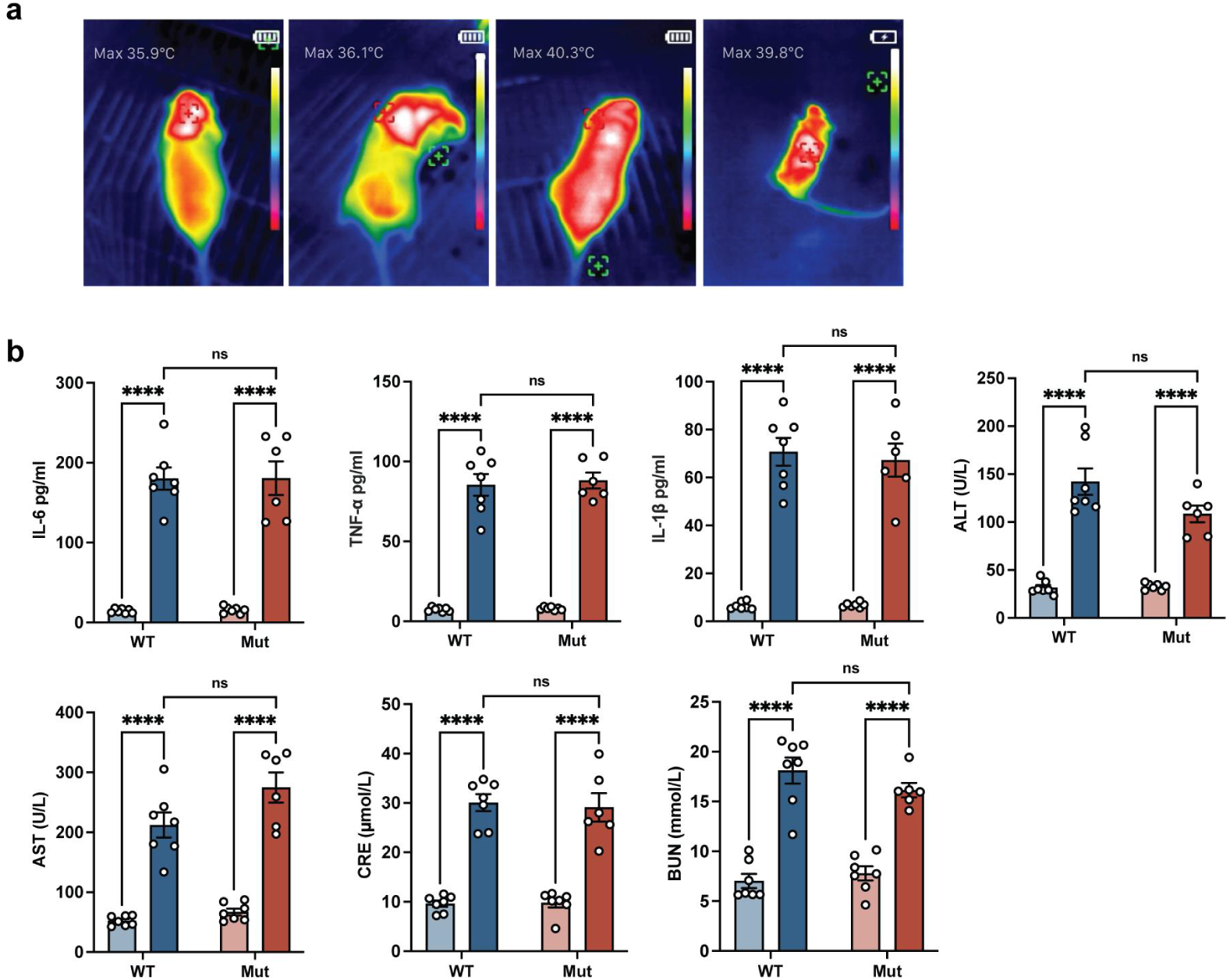
Body-surface temperature and systemic injury markers in the heat-stroke model. Mice and groups are as in Fig. 4: each group began with seven mice, and one Mut+HS mouse died, giving n = 6 for that group and n = 7 for all others. **a,** Representative infrared thermal images taken approximately 30 min into the exposure, at which point mean core temperature was within 0.3 K of its peak, from left to right WT+Sham, Mut+Sham, WT+HS and Mut+HS. The maximum body-surface temperature in each field is the thermal camera’s own annotation and is given in degrees Celsius. These are single animals, and body-surface temperature is not a quantitative measure of thermal load; core temperature for all animals is in Fig. 4b. **b,** Serum IL-6, TNF-α, IL-1β, ALT, AST, creatinine (CRE) and blood urea nitrogen (BUN). Bars are mean ± s.e.m.; points are individual mice. Two-way ANOVA per marker. Within each genotype the sham versus heat-stroke comparison is marked; the genotype × surgery interaction, which is the test of whether the knock-in modifies systemic injury, was corrected across the seven markers by the Holm method. \*\*\*\**P* < 0.0001; ns, not significant.

## Supplementary Tables

**Supplementary Table 1.** Anchor separation of the PHB2 candidate LIR in PHB1-PHB2 structures. Cα-Cα distance between Tyr248 and Leu251, measured from the deposited coordinates of every PHB2 chain in which both residues are modelled, in three cryo-electron microscopy structures covering the closed and open states of the complex.

| pdb | resolution_Å | conformation | chain | ca_ca_Å |
| --- | --- | --- | --- | --- |
| 9O9Z | 2.4 | closed | a | 5.013 |
| 9O9Z | 2.4 | closed | b | 5.016 |
| 9O9Z | 2.4 | closed | c | 5.01 |
| 9O9Z | 2.4 | closed | d | 5.057 |
| 9O9Z | 2.4 | closed | e | 5.025 |
| 9O9Z | 2.4 | closed | f | 5.052 |
| 9O9Z | 2.4 | closed | g | 5.076 |
| 9O9Z | 2.4 | closed | h | 5.06 |
| 9O9Z | 2.4 | closed | i | 5.039 |
| 9O9Z | 2.4 | closed | j | 5.068 |
| 9O9Z | 2.4 | closed | k | 5.07 |
| 9OA0 | 3.1 | open | a | 5.026 |
| 9OA0 | 3.1 | open | b | 5.036 |
| 9OA0 | 3.1 | open | c | 5.023 |
| 9OA0 | 3.1 | open | d | 5.036 |
| 9OA0 | 3.1 | open | e | 5.038 |
| 9OA0 | 3.1 | open | f | 5.036 |
| 9OA0 | 3.1 | open | g | 5.029 |
| 9OA0 | 3.1 | open | h | 5.025 |
| 9OA0 | 3.1 | open | i | 5.016 |
| 9OA0 | 3.1 | open | j | 4.943 |
| 9OA0 | 3.1 | open | k | 5.054 |
| 9UNL | 2.8 | open | B | 5.161 |
| 9UNL | 2.8 | open | D | 5.137 |
| 9UNL | 2.8 | open | F | 5.116 |
| 9UNL | 2.8 | open | H | 5.148 |
| 9UNL | 2.8 | open | J | 5.22 |
| 9UNL | 2.8 | open | L | 5.266 |
| 9UNL | 2.8 | open | N | 5.085 |
| 9UNL | 2.8 | open | P | 5.172 |
| 9UNL | 2.8 | open | R | 5.013 |
| 9UNL | 2.8 | open | T | 5.099 |
| 9UNL | 2.8 | open | V | 5.126 |

**Supplementary Table 2.** Anchor separation in ATG8-family complexes with a canonical LIR. Cα-Cα distance between the aromatic ([W/F/Y]) and aliphatic ([L/I/V]) anchors of the [W/F/Y]-x-x-[L/I/V] motif, measured from the deposited coordinates of seven human complexes. Both anchors were required to contact the receptor (minimum heavy-atom distance < 4.5 Å), so that the motif is engaged in the groove rather than merely present in the sequence. The UBA5 peptide bound to GABARAPL2 (PDB 6H8C) was excluded: its binding element is a UFIM with i, i+4 anchor spacing rather than the i, i+3 of a canonical LIR.

| pdb | resolution_Å | receptor | ligand | anchor_i | anchor_i3 | ca_ca_Å |
| --- | --- | --- | --- | --- | --- | --- |
| 3WIM | 2 | GABARAP | WDFY3/ALFY | Phe3346 | Val3349 | 8.63 |
| 8Q6Q | 1.8 | GABARAPL2 | IRGQ | Phe186 | Leu189 | 8.99 |
| 5LXI | 2.3 | GABARAPL1 | ATG4B | Phe388 | Leu391 | 9.01 |
| 5CX3 | 1.8 | LC3A | FYCO1 | Phe1280 | Ile1283 | 9.12 |
| 5YIR | 1.9 | GABARAP | Ankyrin-2 | Trp1989 | Val1992 | 9.12 |
| 5D94 | 2.2 | LC3B | FYCO1 | Phe1280 | Ile1283 | 9.3 |
| 2ZJD | 1.56 | LC3B | p62/SQSTM1 | Trp340 | Leu343 | 9.63 |

**Supplementary Table 3.**
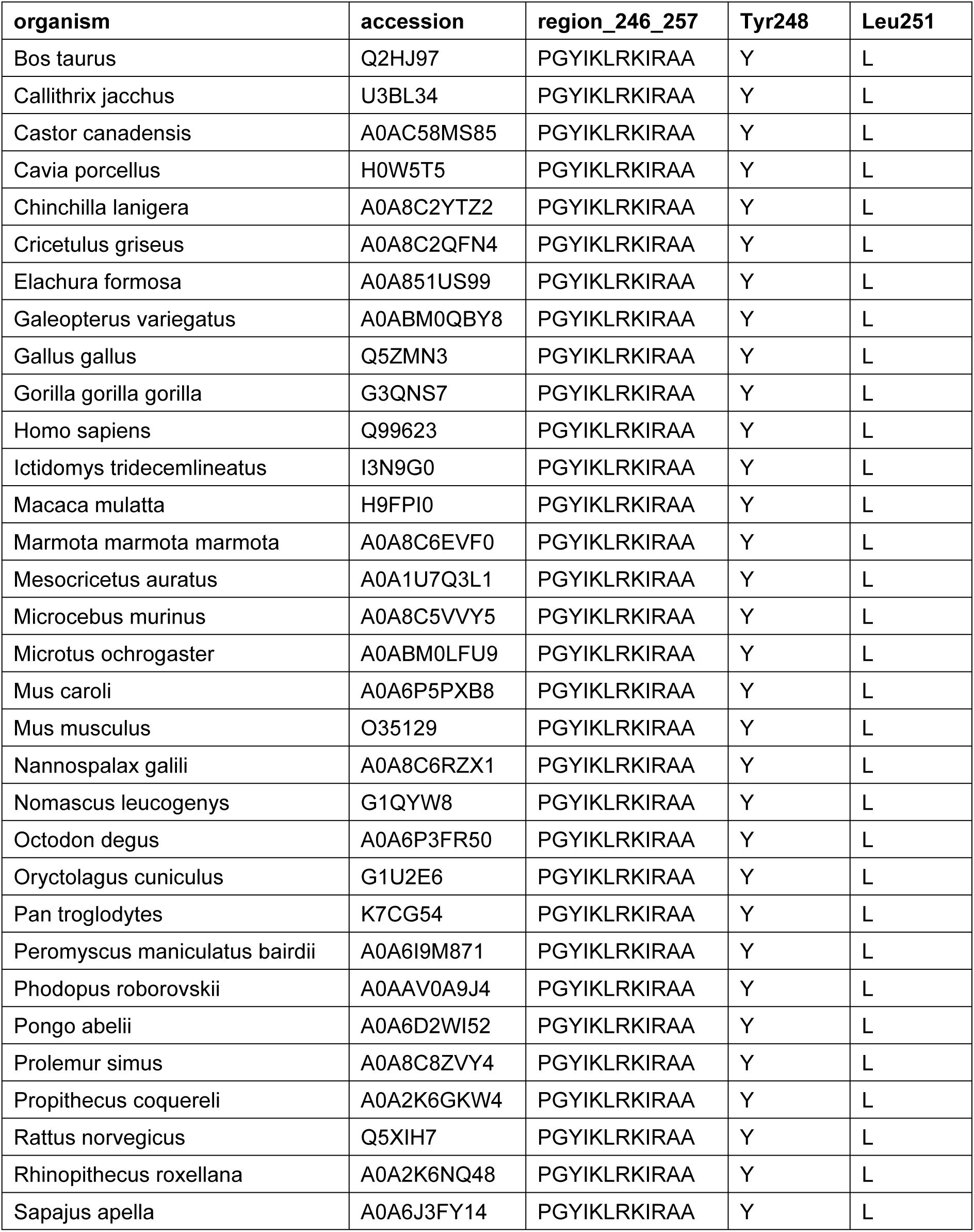
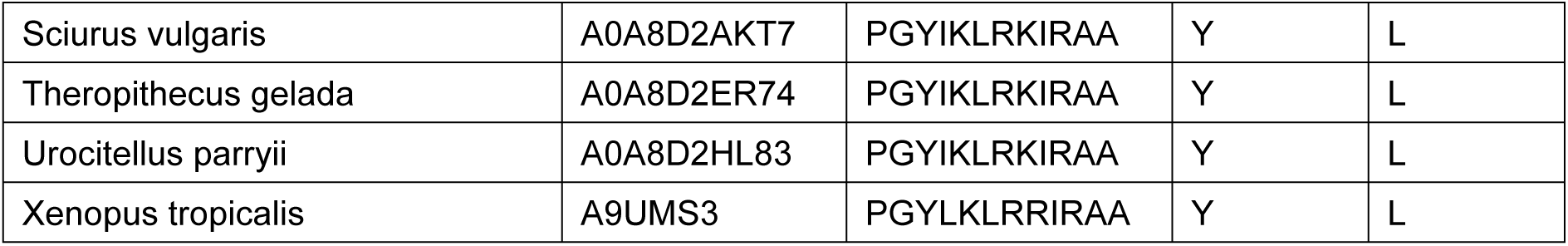
Conservation of PHB2 residues 246-257 across vertebrate orthologues. PHB2 sequences were retrieved from UniProtKB (gene:PHB2, taxonomy_id:7742 Vertebrata, together with the reviewed PHB2 set). In each orthologue the segment was located by matching the 20 residues that precede it in human (226-245) and taking the following 12, rather than by residue number, which differs between species.

**Supplementary Table 4.**
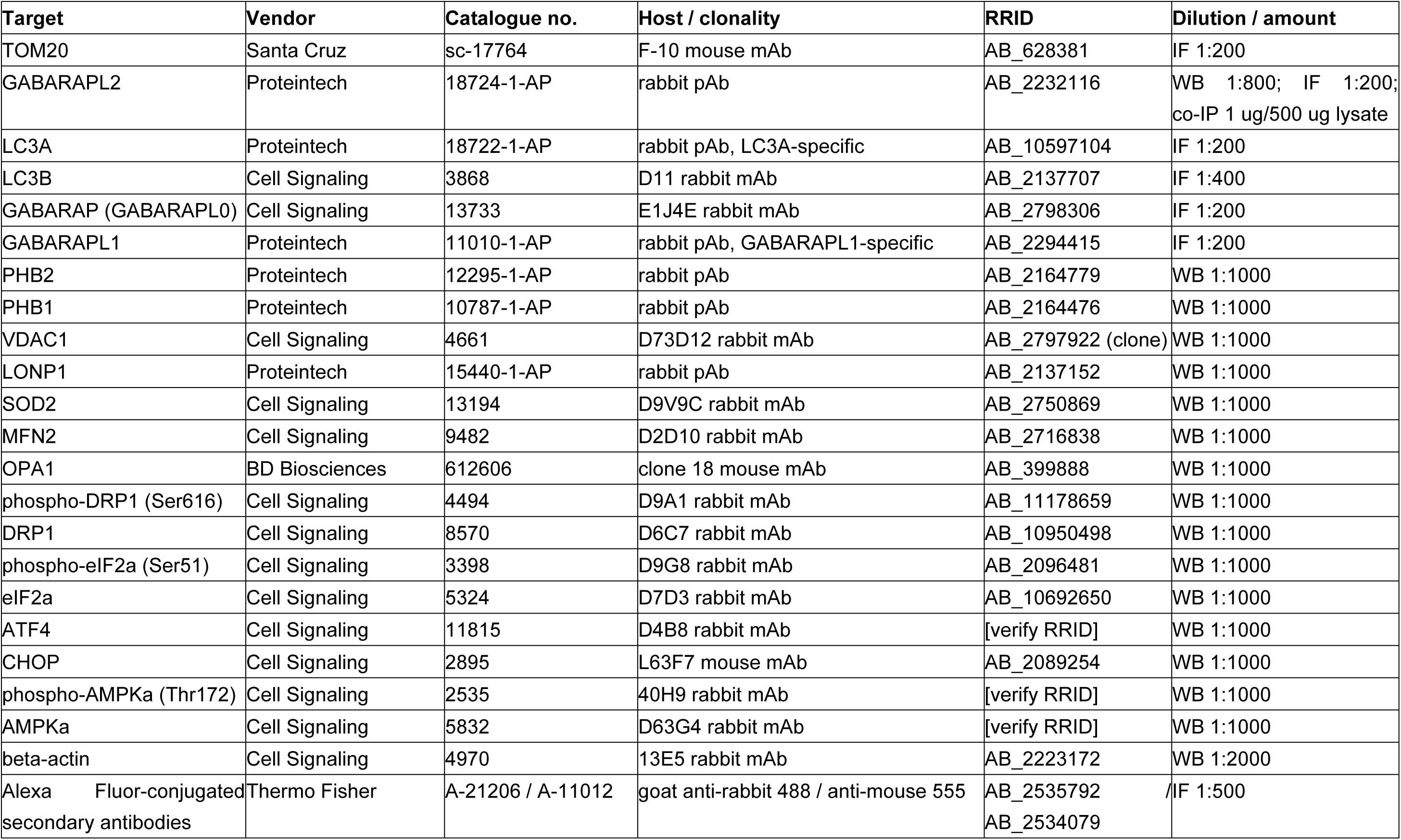
Antibodies used in this study.

**Supplementary Fig. 1.**
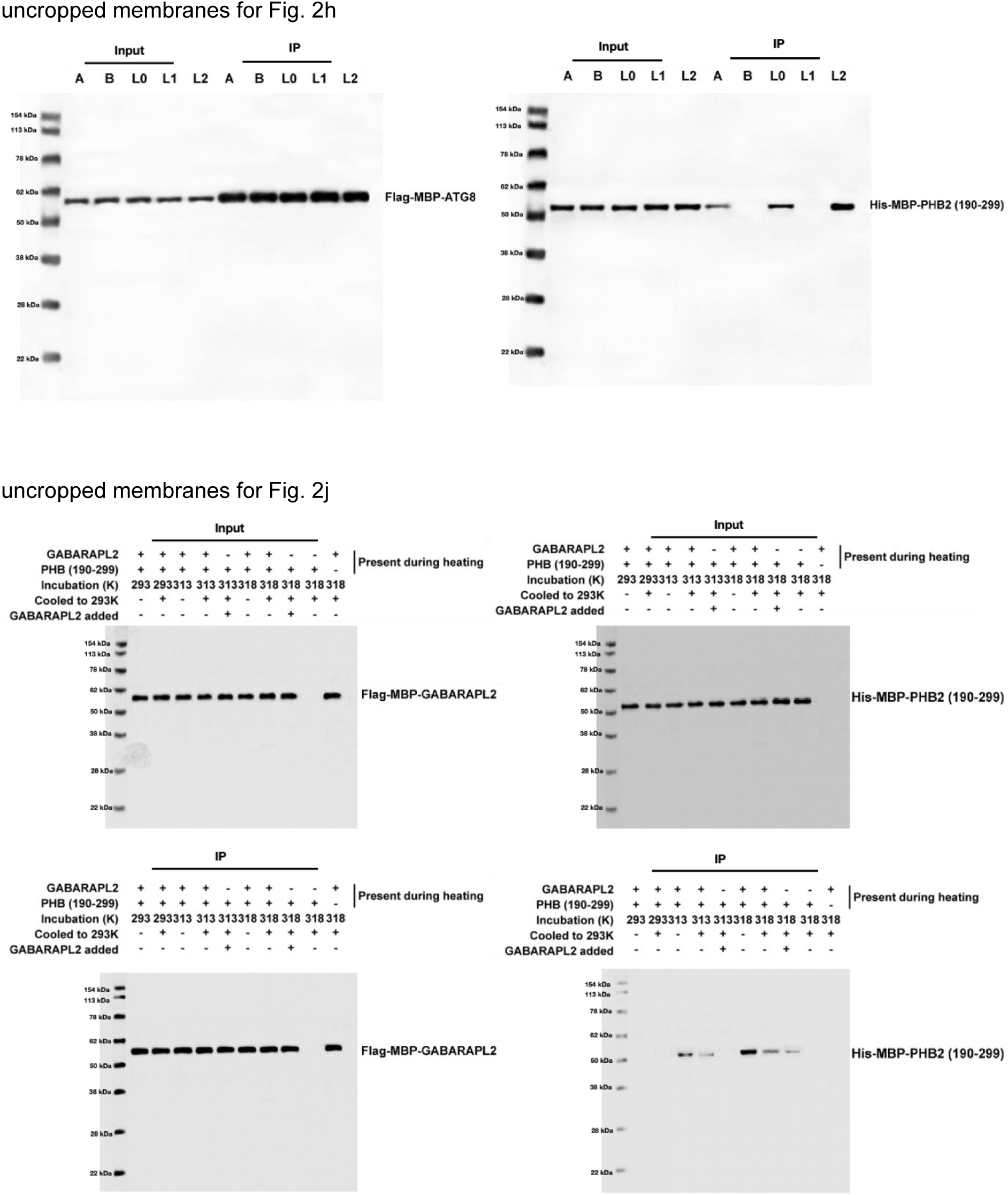
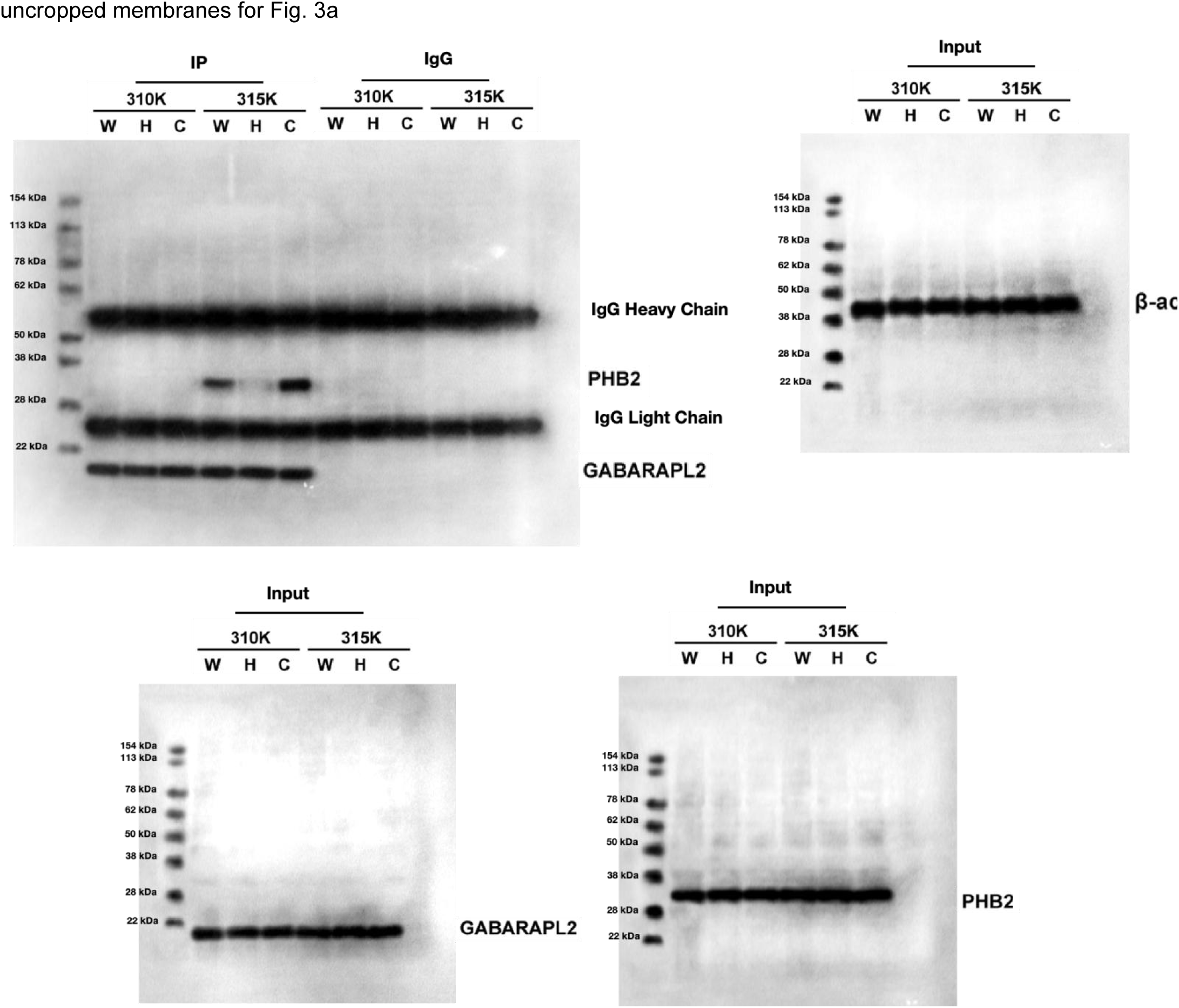
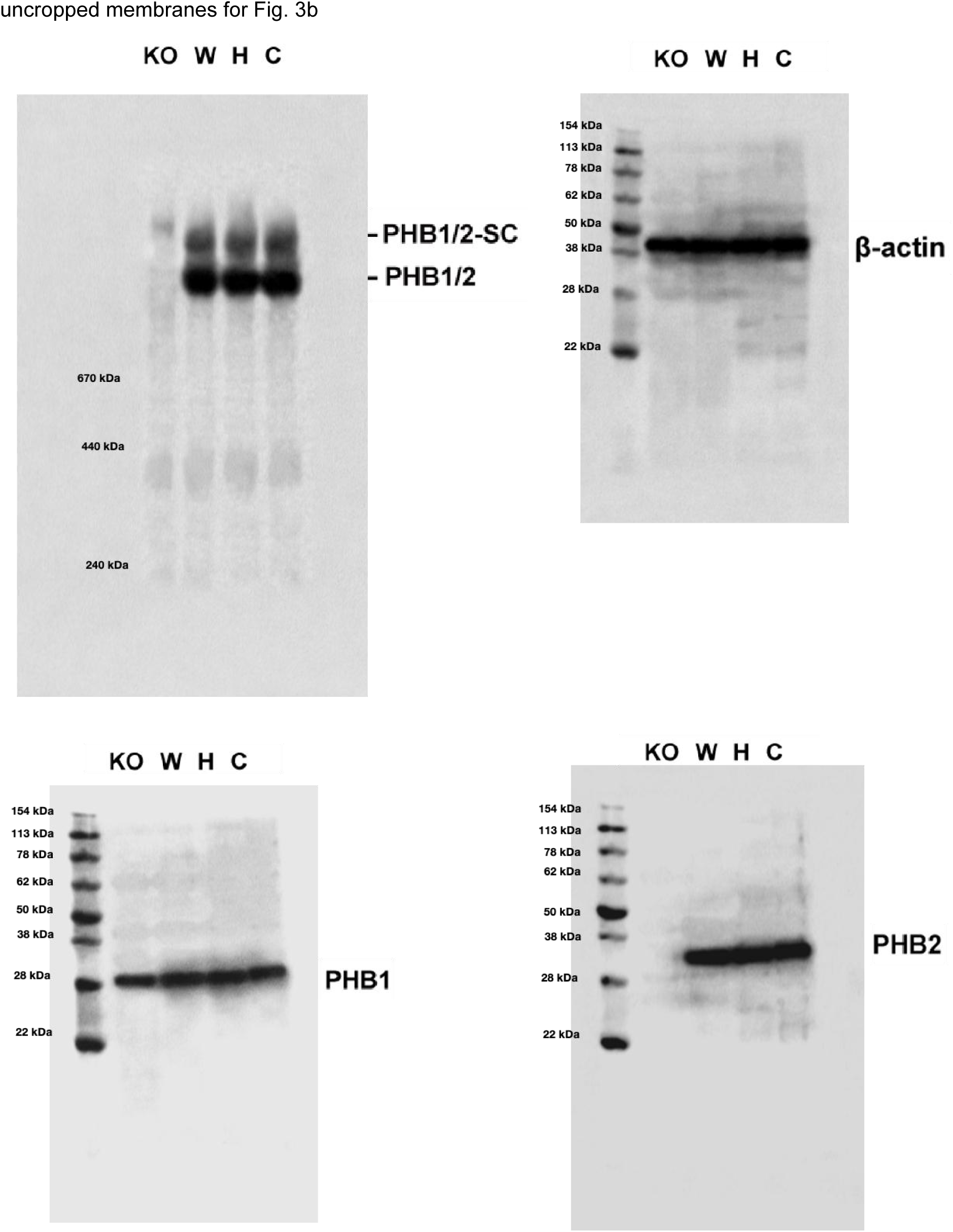
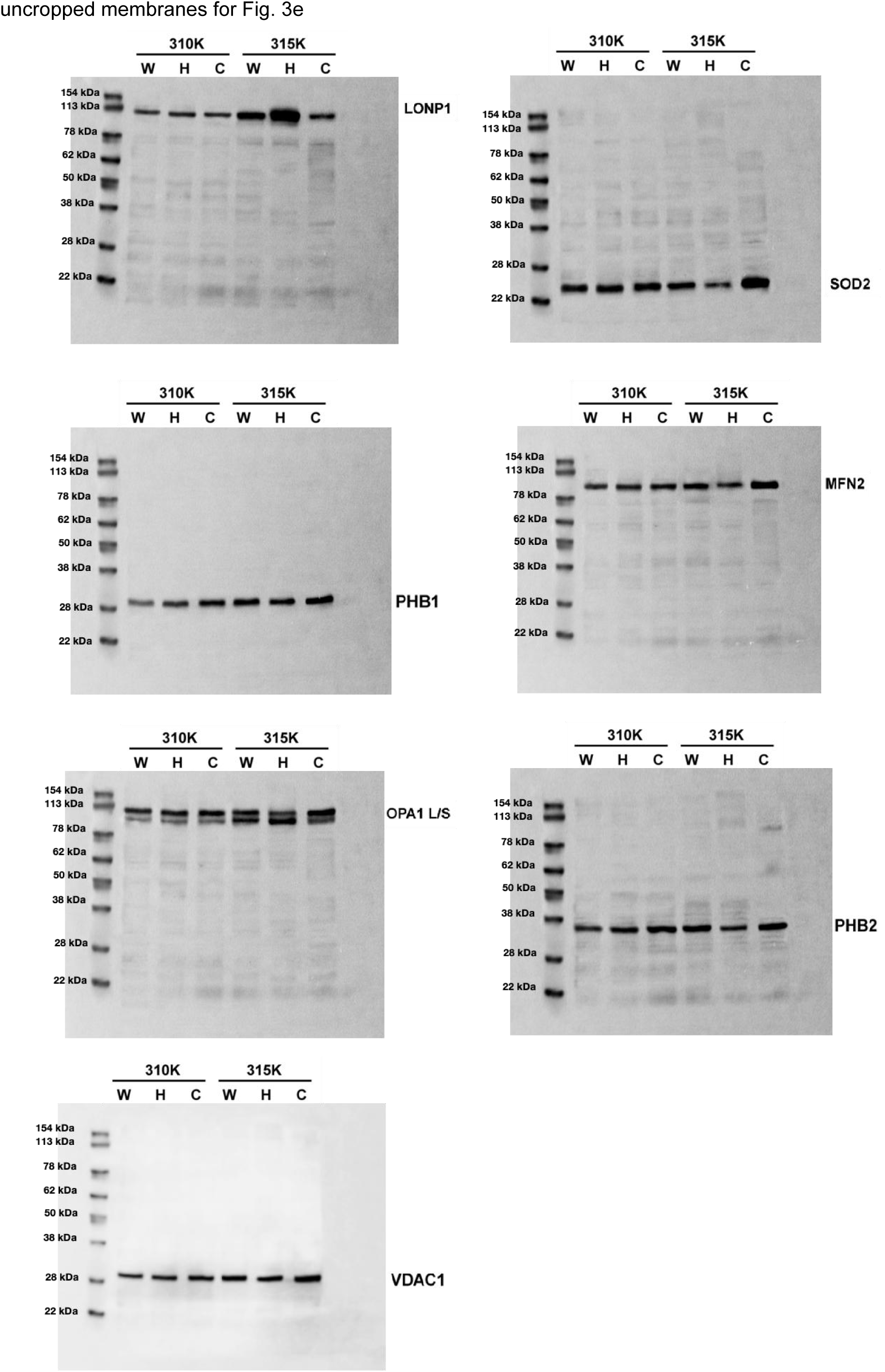
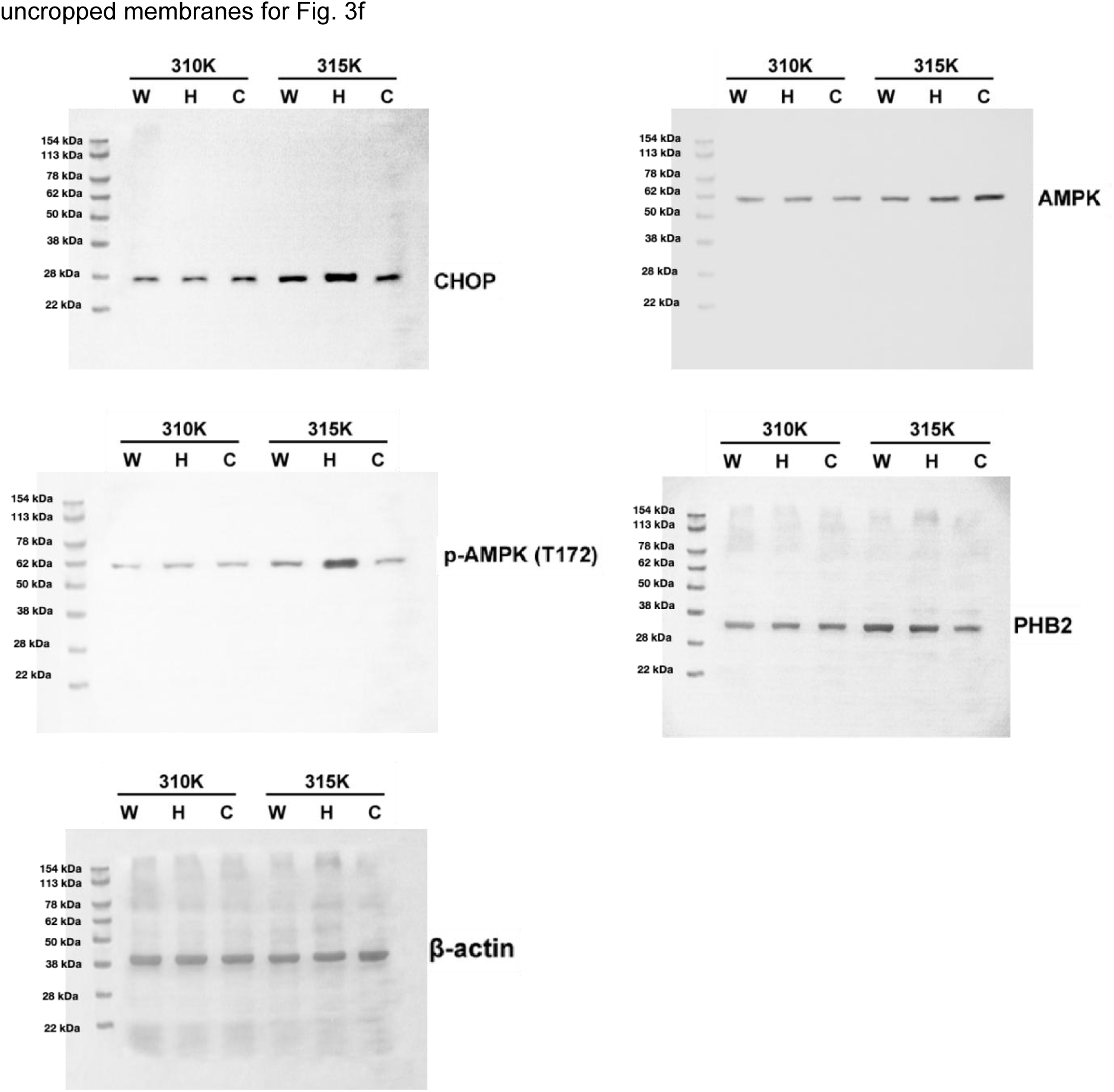
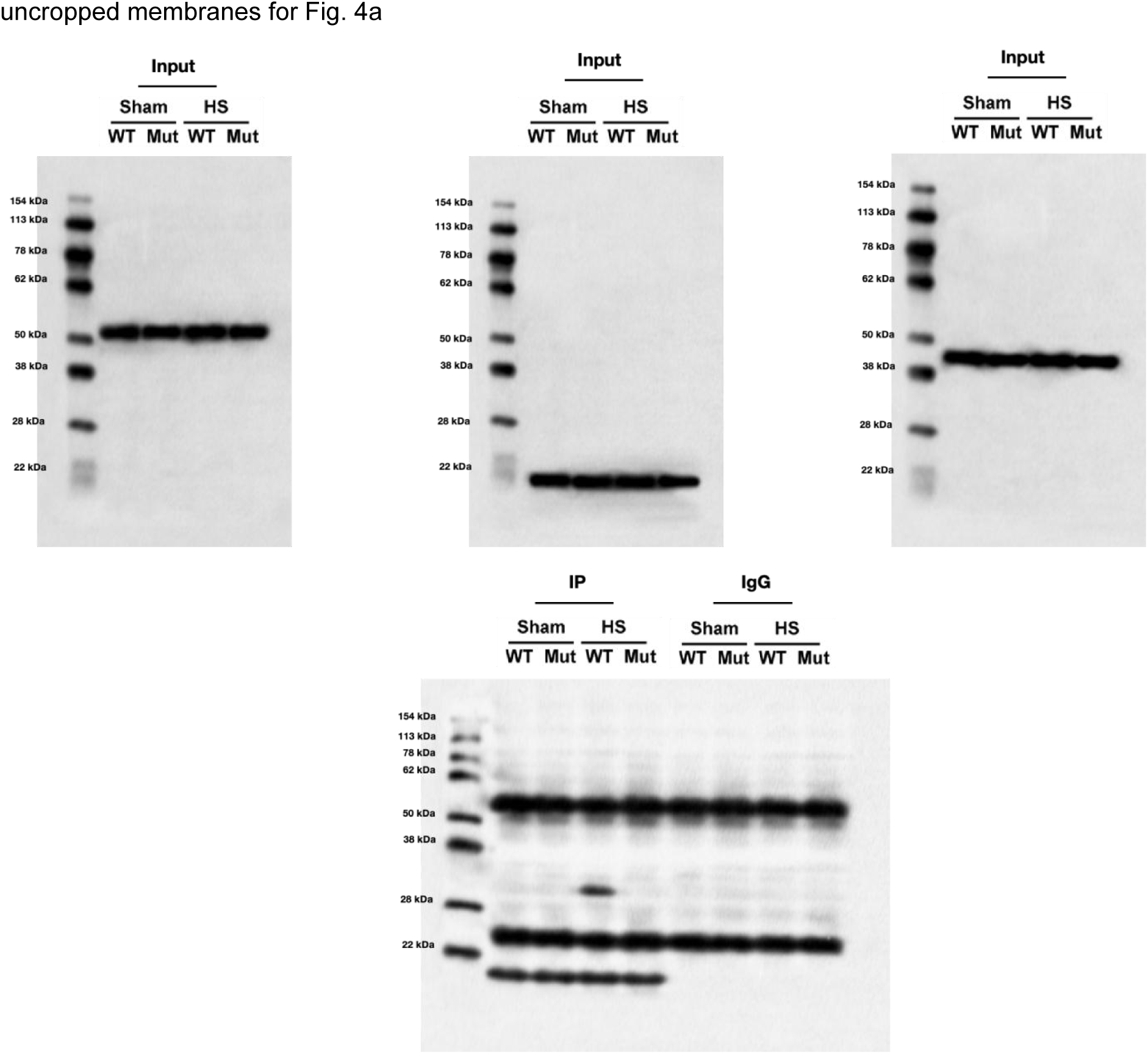
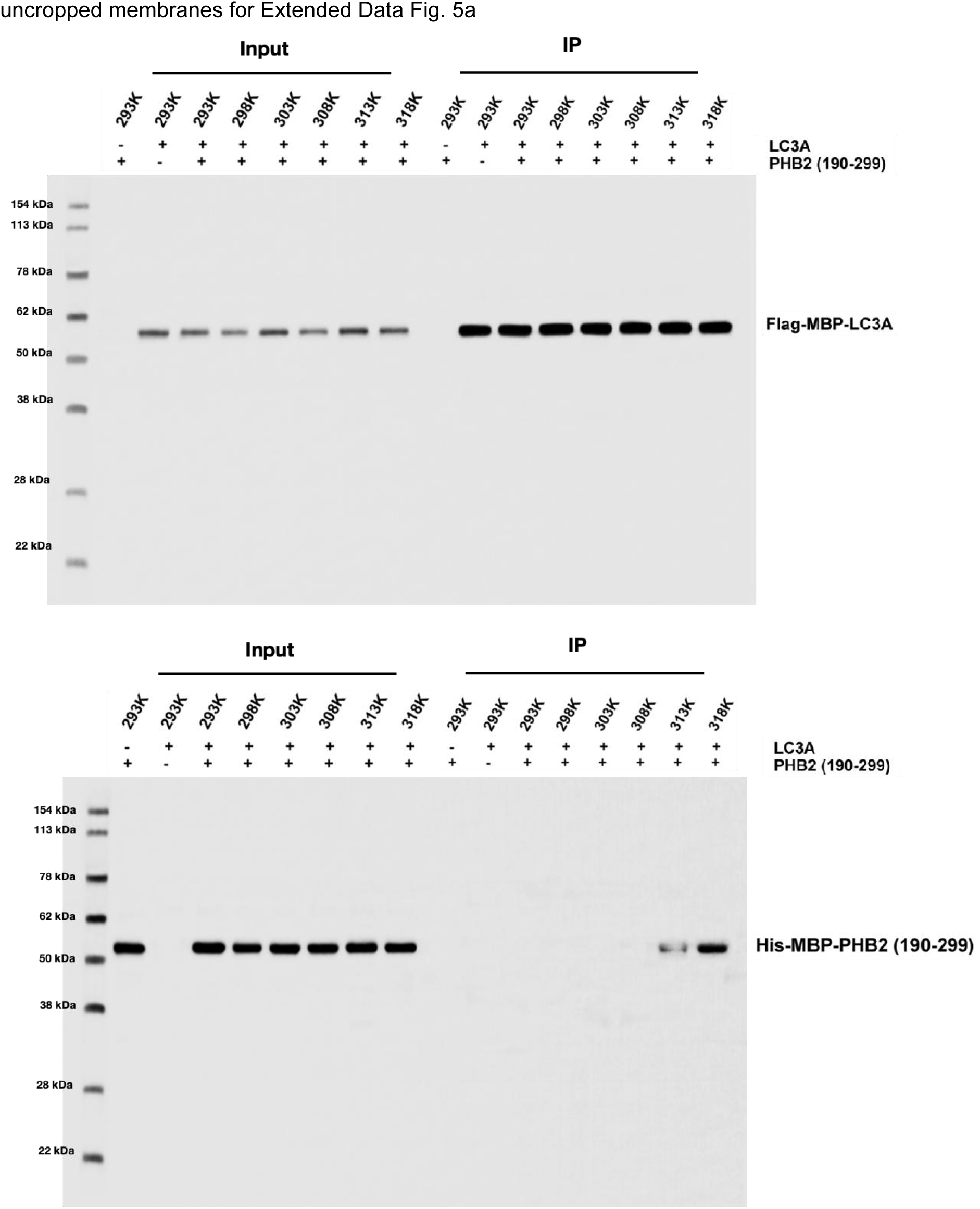
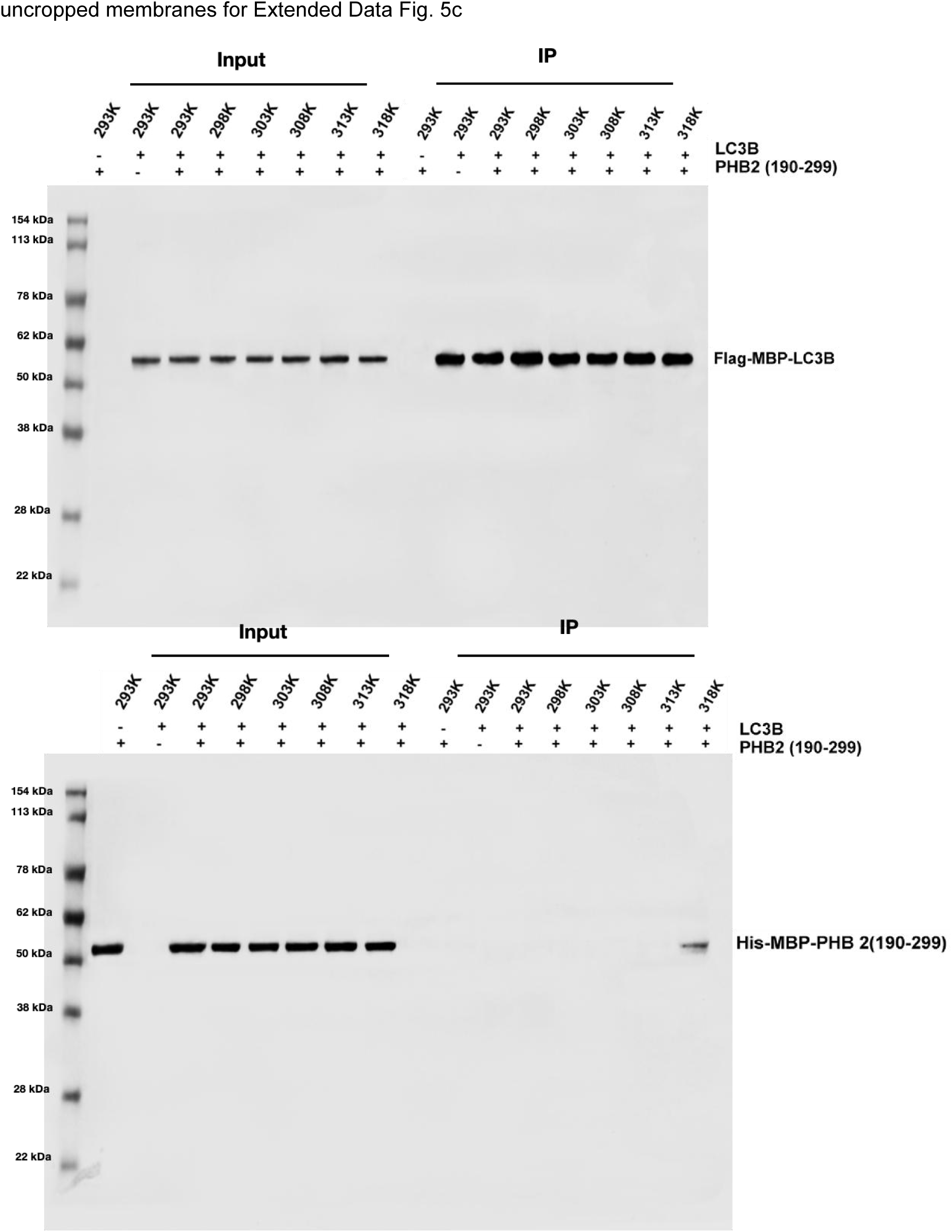
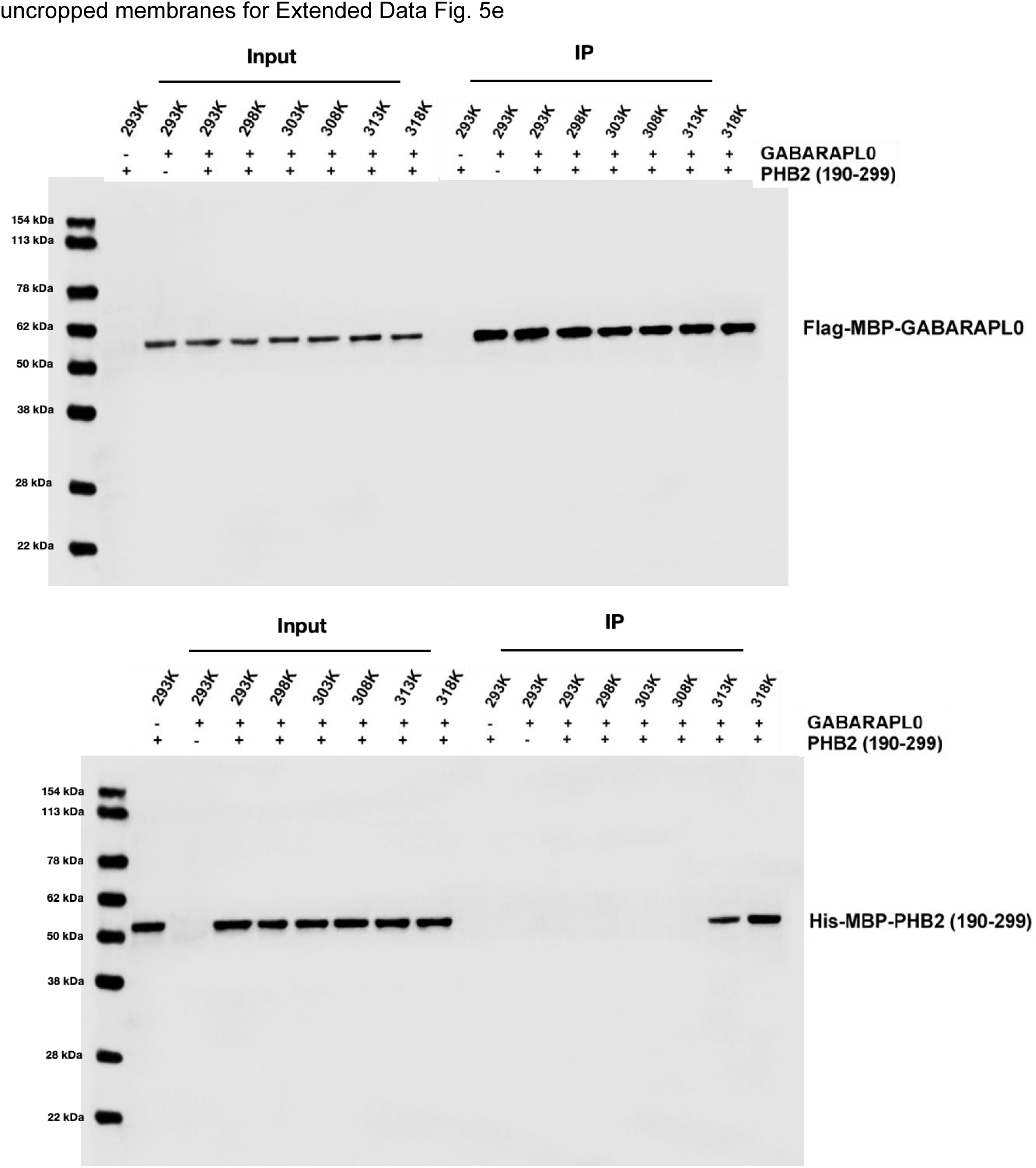
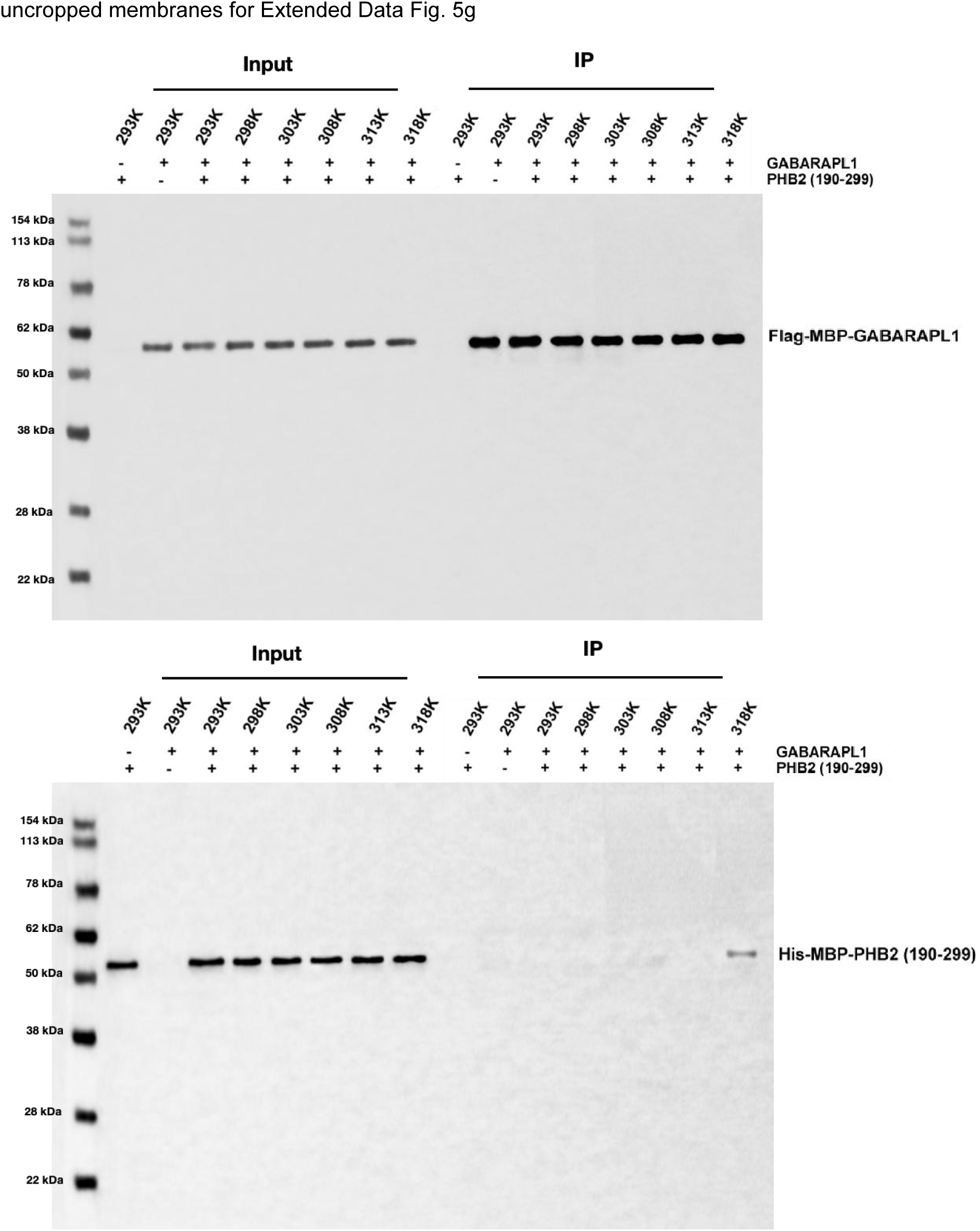
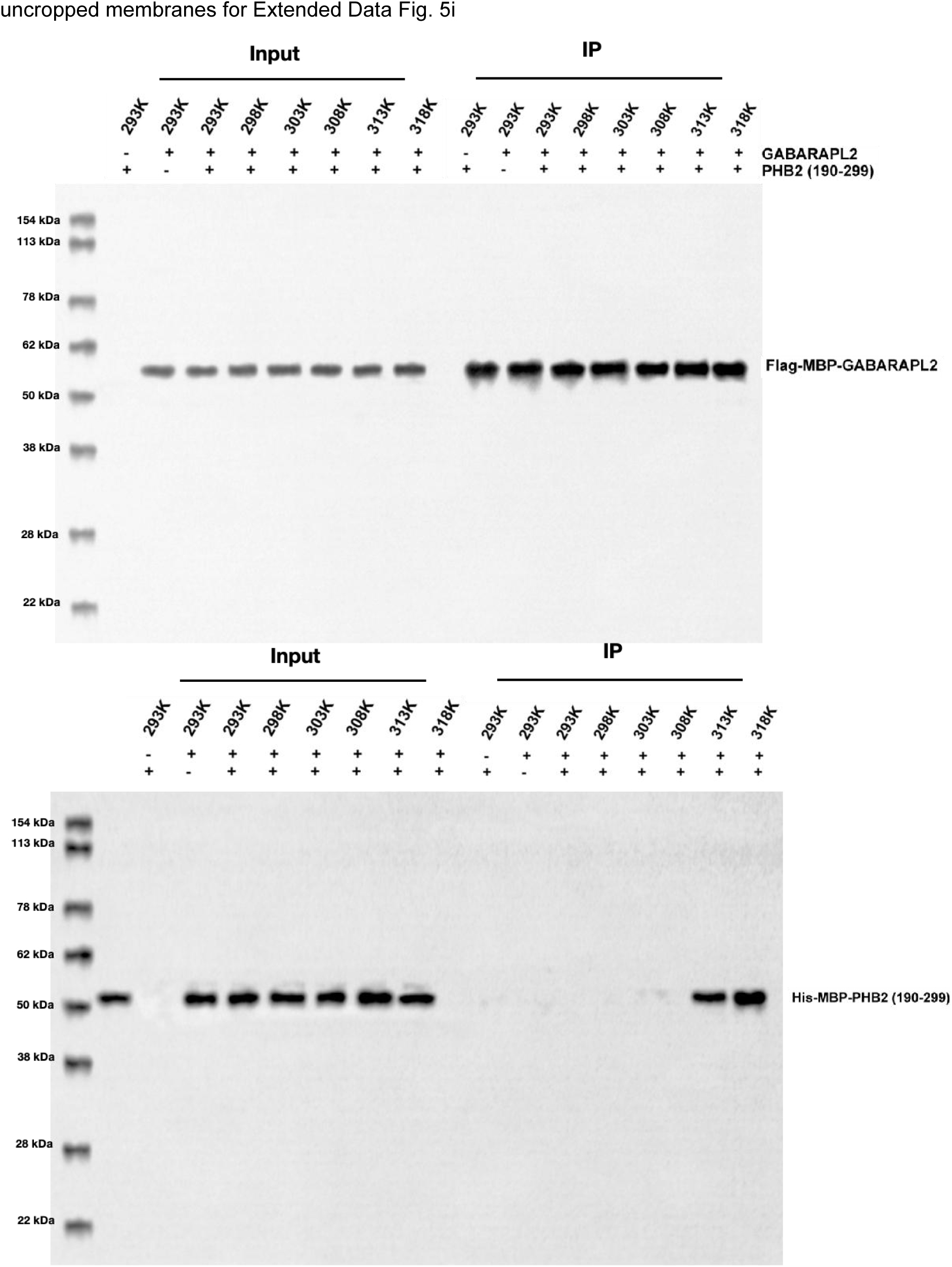
Uncropped immunoblots. Full membranes for every immunoblot shown in the main and Extended Data figures.

## References

1 Mezzacappo, N. F. et al. Uncovering the time-temperature thresholds of in vitro mitochondrial bioenergetics dysfunction under hyperthermic stress. Int J Hyperthermia 42, 2560017 (2025). 10.1080/02656736.2025.2560017

2 Zukiene, R., Nauciene, Z., Ciapaite, J. & Mildaziene, V. Acute temperature resistance threshold in heart mitochondria: Febrile temperature activates function but exceeding it collapses the membrane barrier. Int J Hyperthermia 26, 56–66 (2010). 10.3109/02656730903262140

3 Xia, R. et al. The pathogenesis and therapeutic strategies of heat stroke-induced myocardial injury. Front Pharmacol 14, 1286556 (2023). 10.3389/fphar.2023.1286556

4 Zhuang, Y., Zhuang, X. H., Zhang, X. Y., Wang, D. C. & Yang, Y. Mechanisms and Intervention Strategies for Heat Stroke-Associated Myocardial Dysfunction: A Narrative Review. West J Emerg Med 27, 526–533 (2026). 10.5811/westjem.53045

5 Bao, Q., Wang, Z., Wang, J. & Ruan, Y. Epidemiology of Ischemic Heart Disease Burden Attributable to High Temperature in Asia From GBD 2021. JACC Asia 5, 528–540 (2025). 10.1016/j.jacasi.2024.12.012

6 Katznelson, E. Temperature-Related Ischemic Heart Disease in Asia: Addressing the Growing Burden. JACC Asia 5, 541–542 (2025). 10.1016/j.jacasi.2025.02.011

7 Liu, L., Sakakibara, K., Chen, Q. & Okamoto, K. Receptor-mediated mitophagy in yeast and mammalian systems. Cell Res 24, 787–795 (2014). 10.1038/cr.2014.75

8 Liu, L. et al. Mitochondrial outer-membrane protein FUNDC1 mediates hypoxia-induced mitophagy in mammalian cells. Nat Cell Biol 14, 177–185 (2012). 10.1038/ncb2422

9 Zhu, Y. et al. Modulation of serines 17 and 24 in the LC3-interacting region of Bnip3 determines pro-survival mitophagy versus apoptosis. J Biol Chem 288, 1099–1113 (2013). 10.1074/jbc.M112.399345

10 Novak, I. et al. Nix is a selective autophagy receptor for mitochondrial clearance. EMBO Rep 11, 45–51 (2010). 10.1038/embor.2009.256

11 Jacomin, A. C., Samavedam, S., Promponas, V. & Nezis, I. P. iLIR database: A web resource for LIR motif-containing proteins in eukaryotes. Autophagy 12, 1945–1953 (2016). 10.1080/15548627.2016.1207016

12 Utichi, M. et al. Decoding phospho-regulation and flanking regions in autophagy-associated short linear motifs. Commun Biol 8, 1255 (2025). 10.1038/s42003-025-08399-9

13 Stuke, J. F. M. & Hummer, G. AlphaFold2 SLiM screen for LC3-LIR interactions in autophagy. Autophagy 21, 2192–2212 (2025). 10.1080/15548627.2025.2493999

14 Wei, Y., Chiang, W. C., Sumpter, R., Jr., Mishra, P. & Levine, B. Prohibitin 2 Is an Inner Mitochondrial Membrane Mitophagy Receptor. Cell 168, 224–238.e210 (2017). 10.1016/j.cell.2016.11.042

15 Lange, F. et al. In situ architecture of the human prohibitin complex. Nat Cell Biol 27, 633–640 (2025). 10.1038/s41556-025-01620-1

16 Gao, J. et al. Structures of human organellar SPFH protein complexes. Nat Commun 16, 10064 (2025). 10.1038/s41467-025-65078-3

17 Hong, S. et al. Cryo-EM structure of the prohibitin complex in open conformation. Proc Natl Acad Sci U S A 122, e2512430122 (2025). 10.1073/pnas.2512430122

18 Herhaus, L. et al. IRGQ-mediated autophagy in MHC class I quality control promotes tumor immune evasion. Cell 187, 7285–7302.e7229 (2024). 10.1016/j.cell.2024.09.048

19 Eisenberg, D., Weiss, R. M. & Terwilliger, T. C. The hydrophobic moment detects periodicity in protein hydrophobicity. Proc Natl Acad Sci U S A 81, 140–144 (1984). 10.1073/pnas.81.1.140

20 Wang, L., Friesner, R. A. & Berne, B. J. Replica exchange with solute scaling: a more efficient version of replica exchange with solute tempering (REST2). J Phys Chem B 115, 9431–9438 (2011). 10.1021/jp204407d

21 Pace, C. N. & Scholtz, J. M. A helix propensity scale based on experimental studies of peptides and proteins. Biophys J 75, 422–427 (1998). 10.1016/s0006-3495(98)77529-0

22 Abramson, J. et al. Accurate structure prediction of biomolecular interactions with AlphaFold 3. Nature 630, 493–500 (2024). 10.1038/s41586-024-07487-w

23 Rogov, V. V. et al. Atg8 family proteins, LIR/AIM motifs and other interaction modes. Autophagy Rep 2 (2023). 10.1080/27694127.2023.2188523

## References

24 Labesse, G., Colloc’h, N., Pothier, J. & Mornon, J. P. P-SEA: a new efficient assignment of secondary structure from C alpha trace of proteins. Comput Appl Biosci 13, 291–295 (1997). 10.1093/bioinformatics/13.3.291

25 Schindelin, J. et al. Fiji: an open-source platform for biological-image analysis. Nat Methods 9, 676–682 (2012). 10.1038/nmeth.2019

26 Costes, S. V. et al. Automatic and quantitative measurement of protein-protein colocalization in live cells. Biophys J 86, 3993–4003 (2004). 10.1529/biophysj.103.038422

27 Manders, E. M. M., Verbeek, F. J. & Aten, J. A. Measurement of co-localization of objects in dual-colour confocal images. J Microsc 169, 375–382 (1993). 10.1111/j.1365-2818.1993.tb03313.x

28 Wittig, I., Braun, H. P. & Schägger, H. Blue native PAGE. Nat Protoc 1, 418–428 (2006). 10.1038/nprot.2006.62

29 UniProt: the Universal Protein Knowledgebase in 2025. Nucleic Acids Res 53, D609–d617 (2025). 10.1093/nar/gkae1010

30 Kabsch, W. & Sander, C. Dictionary of protein secondary structure: pattern recognition of hydrogen-bonded and geometrical features. Biopolymers 22, 2577–2637 (1983). 10.1002/bip.360221211

31 Genheden, S. & Ryde, U. The MM/PBSA and MM/GBSA methods to estimate ligand-binding affinities. Expert Opin Drug Discov 10, 449–461 (2015). 10.1517/17460441.2015.1032936

32 Huang, J. et al. CHARMM36m: an improved force field for folded and intrinsically disordered proteins. Nat Methods 14, 71–73 (2017). 10.1038/nmeth.4067

33 Tribello, G. A., Bonomi, M., Branduardi, D., Camilloni, C. & Bussi, G. PLUMED 2: New feathers for an old bird. Computer Physics Communications 185, 604–613 (2014). 10.1016/j.cpc.2013.09.018

34 McGibbon, R. T. et al. MDTraj: A Modern Open Library for the Analysis of Molecular Dynamics Trajectories. Biophys J 109, 1528–1532 (2015). 10.1016/j.bpj.2015.08.015

35 Baker, E. N. & Hubbard, R. E. Hydrogen bonding in globular proteins. Prog Biophys Mol Biol 44, 97–179 (1984). 10.1016/0079-6107(84)90007-5

36 Shrake, A. & Rupley, J. A. Environment and exposure to solvent of protein atoms. Lysozyme and insulin. J Mol Biol 79, 351–371 (1973). 10.1016/0022-2836(73)90011-9

37 Case, D. A. et al. AmberTools. J Chem Inf Model 63, 6183–6191 (2023). 10.1021/acs.jcim.3c01153

38 Case, D. A. et al. Recent Developments in Amber Biomolecular Simulations. J Chem Inf Model 65, 7835–7843 (2025). 10.1021/acs.jcim.5c01063

39 Maier, J. A. et al. ff14SB: Improving the Accuracy of Protein Side Chain and Backbone Parameters from ff99SB. J Chem Theory Comput 11, 3696–3713 (2015). 10.1021/acs.jctc.5b00255

40 Michaud-Agrawal, N., Denning, E. J., Woolf, T. B. & Beckstein, O. MDAnalysis: a toolkit for the analysis of molecular dynamics simulations. J Comput Chem 32, 2319–2327 (2011). 10.1002/jcc.21787

41 Holm, S. A Simple Sequentially Rejective Multiple Test Procedure. Scandinavian Journal of Statistics 6, 65–70 (1979).

42 Agah, R. et al. Gene recombination in postmitotic cells. Targeted expression of Cre recombinase provokes cardiac-restricted, site-specific rearrangement in adult ventricular muscle in vivo. J Clin Invest 100, 169–179 (1997). 10.1172/jci119509

